# CryoEM structure of the Vibrio cholerae Type IV competence pilus secretin PilQ

**DOI:** 10.1101/2020.03.03.975896

**Authors:** Sara J. Weaver, Matthew H. Sazinsky, Triana N. Dalia, Ankur B. Dalia, Grant J. Jensen

## Abstract

Natural transformation is the process by which bacteria take up genetic material from their environment and integrate it into their genome by homologous recombination. It represents one mode of horizontal gene transfer and contributes to the spread of traits like antibiotic resistance. In *Vibrio cholerae*, the Type IV competence pilus is thought to facilitate natural transformation by extending from the cell surface, binding to exogenous DNA, and retracting to thread this DNA through the outer membrane secretin, PilQ. A lack of structural information has hindered our understanding of this process, however. Here, we solved the first ever high-resolution structure of a Type IV competence pilus secretin. A functional tagged allele of VcPilQ purified from native *V. cholerae* cells was used to determine the cryoEM structure of the PilQ secretin in amphipol to ∼2.7 Å. This structure highlights for the first time key differences in the architecture of the Type IV competence pilus secretin from the Type II and Type III Secretin System secretins. Based on our cryoEM structure, we designed a series of mutants to interrogate the mechanism of PilQ. These experiments provide insight into the channel that DNA likely traverses to promote the spread of antibiotic resistance via horizontal gene transfer by natural transformation. We prove that it is possible to reduce pilus biogenesis and natural transformation by sealing the gate, suggesting VcPilQ as a new drug target.

## Introduction

Horizontal gene transfer, or the ability of microorganisms to directly share DNA with one another, facilitates rapid evolution, can contribute to the development of antibiotic resistance, promote the spread of virulence factors, and allow bacterial pathogens to rapidly evade host immune response ^1^. A clear understanding of the mechanisms of horizontal gene transfer can aid the development of tools in the fight against antibiotic resistance.

One mechanism of horizontal gene transfer is natural transformation, where a competent bacterium can take up DNA from its environment and then maintain this exogenous genetic material, either as a plasmid or by integrating it into the genome by homologous recombination ^2^. During transformation, DNA is actively brought through the membrane. The requirements for DNA uptake vary with species, but there is evidence for transformation of DNA from a few hundred base pairs up to tens of thousands base pairs ^2–5^. Intergenus transformation can result in the development of mosaic alleles that confer antibiotic resistance, and has been demonstrated in a variety of genera, including *Streptococcus*, *Neisseria*, and *Actinobacter*^6, 7^.

Additionally, natural transformation of large regions of DNA can induce serotype switching in *Vibrio cholerae* ^8, 9^ and in *Streptococcus pneumoniae* ^10^. The serotype of a bacterial strain describes the immunodominant surface antigen that it displays, and adaptive immune responses in human populations are generally serotype specific. This has major public health implications, as serotype switching can help bacteria evade the immune system. For example, after the introduction of a new vaccine against a serotype of *S. pneumoniae*, epidemiologists traced a vaccine-escape serotype to horizontal gene transfer ^10^. Thus, preventing horizontal gene transfer represents a unique approach to mitigate the spread of antibiotic resistance and virulence in bacterial pathogens.

Here, we focus our attention to the structural biology of natural transformation in the gram-negative bacterium *Vibrio cholerae. V. cholerae* is the causative agent of the diarrheal disease cholera ^11^. Since 1817, cholera has spread globally in seven pandemics that each feature strains of distinct characteristics. Natural transformation in *V. cholerae* is tightly regulated and induced when these bacteria are grown on chitin in their aquatic environment ^12^. Chitin, a biopolymer found in the exoskeletons of crustaceans, induces transcription of the chitin regulon and expression of the Type IV competence pilus machinery ^13^. The Type IV competence pilus machinery requires four elements: an inner membrane pilus assembly complex, cytoplasmic motors to extend and retract the pilus, an outer membrane pore (the secretin), and the pilus itself ^14^. The Type IV competence pilus facilitates environmental DNA uptake by extending and retracting from the cell surface through a large, outer membrane secretin pore called PilQ ^15^. To mediate DNA uptake the pilus must translocate DNA across the membrane through the PilQ secretin. Thus, the PilQ secretin represents a potential novel target to thwart this mechanism of horizontal gene transfer.

The Type IV competence secretin is a member of the bacterial Secretin superfamily ^16, 17^. Secretins are found in the Type II Secretion System (T2SS), the T3SS, the Type IV pilus machine, and filamentous phage ^18^. Bacterial secretins are united by a common C-terminal secretin domain that oligomerizes to form a large pore in the outer membrane ^19^. The N-terminal protein domains form rings in the periplasm and are thought to mediate interactions with other proteins ^20, 21^. While the C-terminal secretin domain is remarkably similar across these secretion systems, the N-terminal region varies (**Figure 1A**). The T2SS secretins have N-terminal protein domains N0 to N3, followed by the secretin domain and then the S-domain. The T3SS secretins lack the N2 domain, while the Type IV competence pilus secretins lack the N1, N2, and S-domains. Additionally, the N-terminus of Type IV competence secretins typically include one or more AMIN domains that interact with the peptidoglycan ^22^. These variations in domain architecture are likely related to the specialization of different secretins ^23^. The T2SS exports periplasmic folded proteins to the extracellular space. The T3SS uses a “needle and syringe” to export cytosolic effector proteins outside of the cell, or directly into another cell. The Type IV competence pilus extends and retracts a filament to take up DNA cargo.

**Figure 1:**
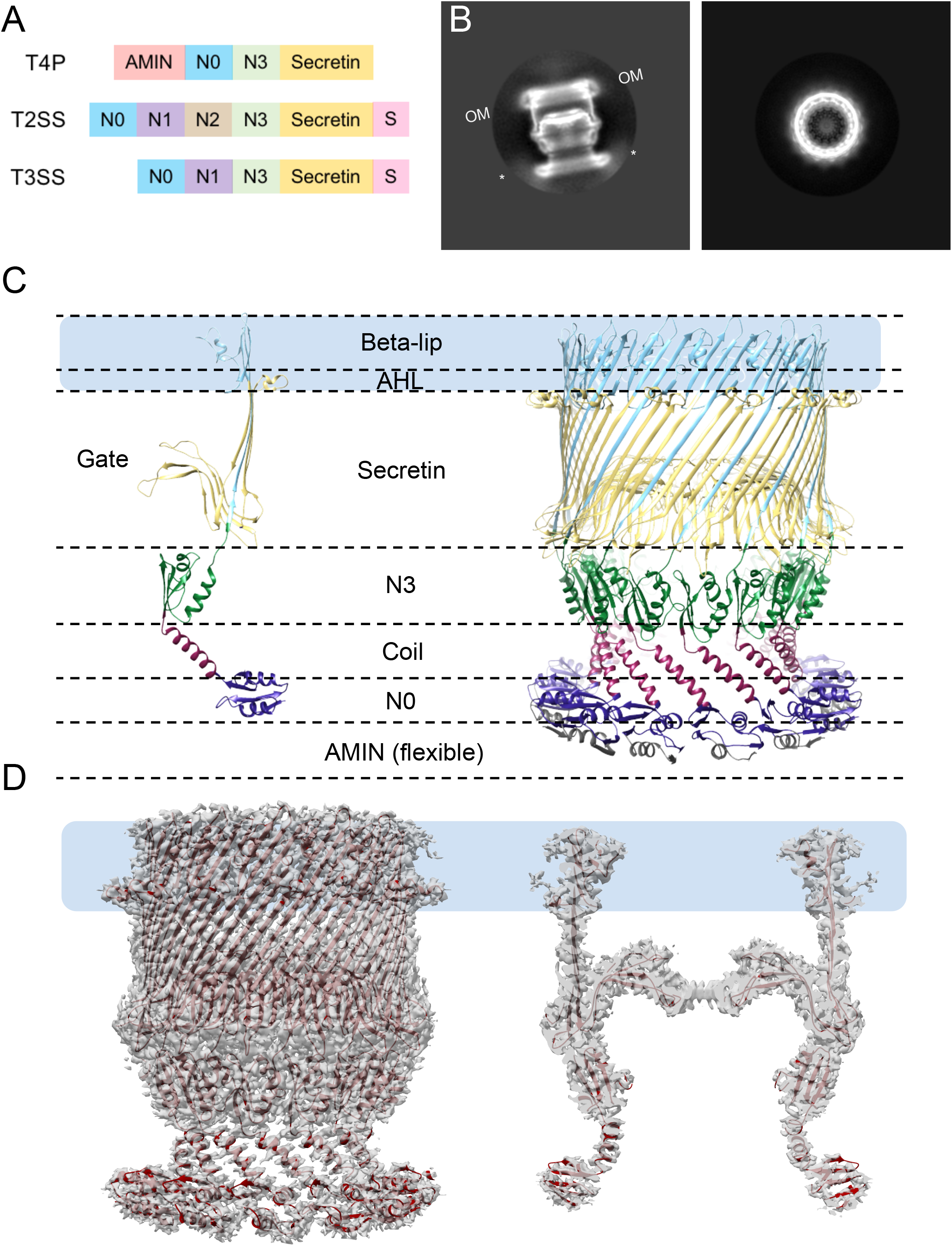
CryoEM structure of the *V. cholerae* Type IV Competence Pilus Secretin PilQ. **(A)** Protein domain organization in the Type IV competence pilus machine, the Type II Secretion System (T2SS) and the Type III Secretion System (T3SS). The AMIN domain (red), the N0 (or STN) domain (blue), the Secretin_N domains N1 (purple), N2 (brown), and N3 (green), the secretin domain (yellow) and the S domain (pink) are shown. **(B)** Example 2D classes show PilQ from the side (left) and from the top (right). The outer membrane is indicated with OM. Asterisks indicate the putative AMIN domains. **(C)** The atomic model of the *V. cholerae* PilQ multimer is shown (right). One chain is shown by itself (left). The outer membrane is represented in a light blue rectangle. Dashed lines represent the different domains of the PilQ structure: AMIN (not shown), N0 (grey/dark blue), coil (purple), N3 (green), Secretin (yellow), Beta-lip (sky blue). The gate is labeled on the monomer. **(D)** Model and cryoEM density of symmetrized PilQ is shown from the side (left) and cut through the center (right).

The structure of the Type IV competence pilus secretin has been examined several times, though no work has provided high-resolution details sufficient to model the passage of DNA or to design inhibitors of this potential drug target. In 2012, a 26 Å structure of *Neisseria meningitidis* PilQ was solved by cryoEM ^20^. In 2016, Koo *et al.* solved the structure of *Pseudomonas aeruginosa* PilQ to 7.4 Å (pink in **Figure 2**). PaPilQ appeared to have 14 peripheral spokes lining the beta barrel that did not demonstrate clear C14 symmetry, so C7 symmetry was assumed. In a 2017 paper, D’Imprima *et al.* described the structure of *T. thermophilus* PilQ to ∼7 Å resolution with reported C13 symmetry (orange in **Figure 2**). D’Imprima *et al.* observed flexibility between the N-terminal domains in TtPilQ that complicated the cryoEM data analysis. Several high resolution cryoEM structures of bacterial T2SS and T3SS secretins have been published, including *E. coli* GspD ^24–26^, *V. cholera* GspD ^24, 26^, *Pseudomonas aeruginosa* XcpQ ^27^, *Klebsiella pneumoniae* PulD ^28^, *A. hydrophila* (pilotin-independent ExeD), *V. vulnificus* (pilotin-dependent EpsD)^29^, and *Salmonella typhimurium* (InvG)^30^. Thus, to date, structural information about the Type IV competence pilus secretin has mainly been inferred from the related, but distinct T2SS and T3SS secretins.

**Figure 2:**
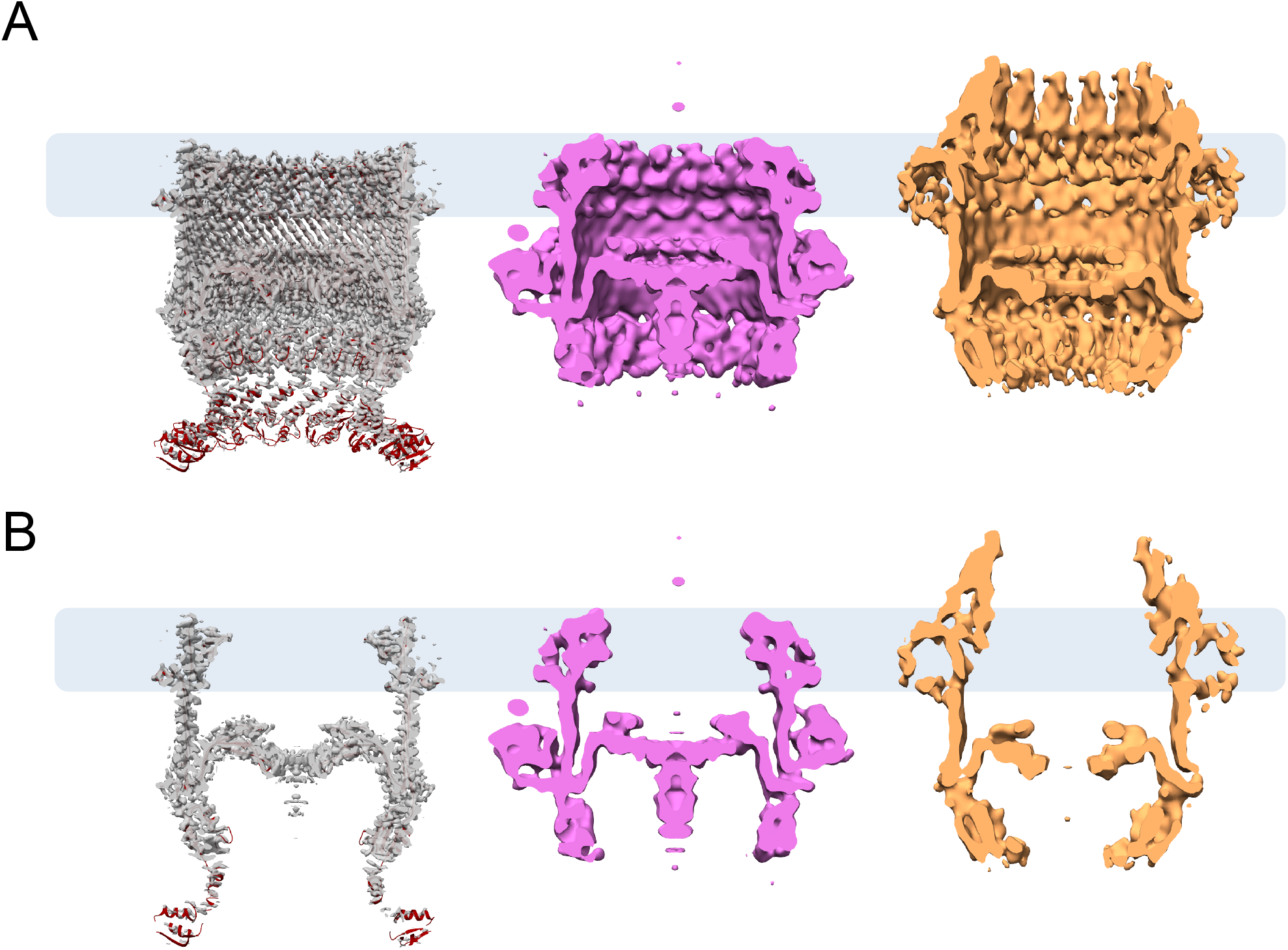
Comparison of VcPilQ to *P. aeruginosa* and *T. thermophilus* PilQ structures. Comparison of cryoEM density of PilQ structures from *V. cholerae* (left, grey)(this study), *Pseudomonas aeruginosa* (magenta, center) (Koo et al. 2016), and *Thermus thermophilus* (orange, right) (D’Imprima et al. 2017). In **(A)**, the inside cavity of the cryoEM densities are shown, while **(B)** highlights the gate region with a cut through the center of the density. The membrane is blue.

Here, we present the structure of the *Vibrio cholerae* Type IV competence pilus secretin PilQ to ∼2.7 Å using a fully functional, His-tagged allele that we expressed and purified from the native bacterium. Our work highlights differences between the Type IV competence pilus, the T2SS, and the T3SS secretins, and emphasizes the need for structures of different secretin family members. In particular we discuss differences and remaining puzzles, including how the pilus could be accommodated within PilQ during natural transformation and what part/s of the secretin if any penetrate the outer membrane. Finally we report structure-inspired designs of cysteine pair mutants that allowed us to reversibly inhibit pilus assembly and natural transformation, presumably by sealing the secretin gate. These experiments support the designation of VcPilQ as a druggable target, and more broadly demonstrate how cysteine pair mutations can be employed to study the activity of bacterial secretins.

## Results

### Purification of *V. cholerae* PilQ in amphipol

To ensure properly-folded and fully-function Type IV competence pilus machinery, we chose to purify PilQ from *Vibrio cholerae* rather than a recombinant system. A chromosomal mutation was made to add a deca-histidine tag to the N-terminus of PilQ. The bacteria retained wild-type levels of natural transformation (**Supplemental Figure 1A**). Previous work demonstrated that similar N-terminal tags allowed for functional Type IV competence pilus activity ^15^.

*V. cholerae* cells expressing PilQ were lysed in the presence of the surfactant n-dodecyl β-D-maltoside (DDM). After affinity purification, PilQ was exchanged into an amphipol environment. Secretins are detergent- and heat-resistant multimers ^31, 32^. SDS page analysis of PilQ in amphipol showed a high molecular weight band near the top of the gel (marked with an asterisk in **Supplemental Figures 1B, D, and E**). Western blotting against the histidine tag on PilQ confirmed the high molecular weight band as PilQ (marked with an asterisk in **Supplemental Figures 1C and F**). The gel electrophoresis also demonstrated some low molecular weight species in the purified sample, some of which are detected on the Western blot and presumed to be PilQ monomer or proteolyzed PilQ species that retained the N-terminal histidine tag. Other bands on the SDS-PAGE not labeled in the Western, likely represent PilQ fragments lacking the N-terminal histidine or contaminant proteins. Regardless, the size difference between these contaminants (<100 kDa) and the PilQ multimer (∼860 kDa) made it easy to distinguish PilQ from the milieu in electron micrographs (**Supplemental Figure 3A)**.

To confirm that full length PilQ was present in our samples (residues 30-571 after the 29 amino acid N-terminal signal peptide is cleaved), gel band analysis with a trypsin digest and mass spectrometry was used to analyze the multimeric PilQ in amphipol from a SDS page gel (band marked with an * in **Supplemental Figure 1D** was analyzed). The results demonstrated 65% sequence coverage, with fragments identified in each domain of the folded protein from residues 50 to 567 (**Supplemental Figure 2**).

### Single particle cryoEM of the Type IV competence pilus secretin PilQ

Here we report the high-resolution structure of the purified Type IV competence pilus secretin VcPilQ by single particle cryoEM (**Figure 1**). The cryoEM data processing steps are summarized as a flow chart in **Supplemental Figure 4** and in Methods.

Briefly, *V. cholerae* PilQ (VcPilQ) was plunge-frozen on Quantifoil Holey Carbon Grids with R2/2 spacing. A Titan Krios operating in a three by three pattern with beam-image shift ^33^ was used to collect 3 movies per hole, resulting in 3,808 movies. The movies were motion corrected and screened for quality, which left 2,510 micrographs.

The cryoSPARC blob picker was used to identify 3,100,353 putative particle location, several micrographs were inspected to adjust the threshold parameters to select the top 252,319 particles for two-dimensional (2D) classification. 2D classification demonstrated a wide variety of orientations, the clear presence of secondary structure (**Supplemental Figure 3B**), and C14 symmetry (**Supplemental Figure 3C**). After the initial 2D classification runs, 66,507 particles were excluded from the dataset and used to generate a nonsense initial model representing contamination. The 185,812 quality particles (‘include’) were used to generate an initial model. A CryoSPARC heterogeneous refinement using the high quality initial model and the contamination initial model was used to further clean the dataset. Several rounds of 3D classification and 3D refinement were used to select 100,543 particles for further analysis in Relion.

After polishing and CTF Refinement, the cryoEM structure (C14 symmetry, overall resolution FSC@0.143 2.7 Å, FSC@0.5 3 Å, **Figure 2**) reached near-atomic resolution and demonstrated isotropic resolution in the X, Y, and Z axes (3DFSC sphericity 0.96) (**Supplemental Figure 5-6**)^34, 35^.

We were able to recognize and model residues 160 to 571 of PilQ (**Figure 1C, Supplemental Movie 1**). Noting that residues 1-29 are cleaved (the N-terminal signal peptide), our mass spectrometric analysis of the purified protein (**Supplemental Figure 2**) demonstrated the presence of residues between 50 and 567, including the AMIN domain (residues 54-125) that is thought to interact with the peptidoglycan in the periplasm ^22^. While the AMIN domain was not resolved in the PilQ structure, hazy density is present in the 2D classification in the region near the PilQ N-terminus where we would expect the AMIN domain (marked with asterisks in **Figure 1B**). Following the AMIN domain, residues 126-159 are predicted by homology modeling to be unstructured (**Supplemental Figure 7**), so they are also likely included in the hazy density. If that region is unstructured, it is a likely source of the conformational heterogeneity observed as a hazy density in **Figure 1B**. Extensive focused classification calculations were unable to improve the resolution in this region (data not shown). Because the cell wall does not have a crystalline structure, and the 14 AMIN domains around the ring must bind to it in different places around the barrel, the AMIN domains are probably not regularly arranged *in situ* either. In *Neisseria meningitidis*, the lipoprotein PilP is thought to bind the AMIN domain and act as a bridge between the inner and outer membrane components of the Type IV pilus machinery ^36^.

### The structure of VcPilQ demonstrates similarities to other secretins

In each PilQ monomer, four beta strands come together to form a beta sheet (**Figure 1C**). Once assembled, PilQ forms a 56-strand beta-barrel. Inside the barrel, two further beta hairpins (beta strand-turn-beta strand each) form a gate. These regions match the topology of other secretin structures (**Figure 3A-C**). Chimera MatchMaker was used to align several published secretin structures to VcPilQ and calculate the root-mean-square deviation (RMSD) between the C-alpha carbons of each pair ^37^ (**Supplemental Figure 8**). In each case, the RMSD calculated over the protein domains present in both structures (N3, secretin, and beta lip domains) was about 1 Å. The outer membrane regions differ significantly, however, in both angle and membrane spanning distance, as discussed in more detail below.

**Figure 3:**
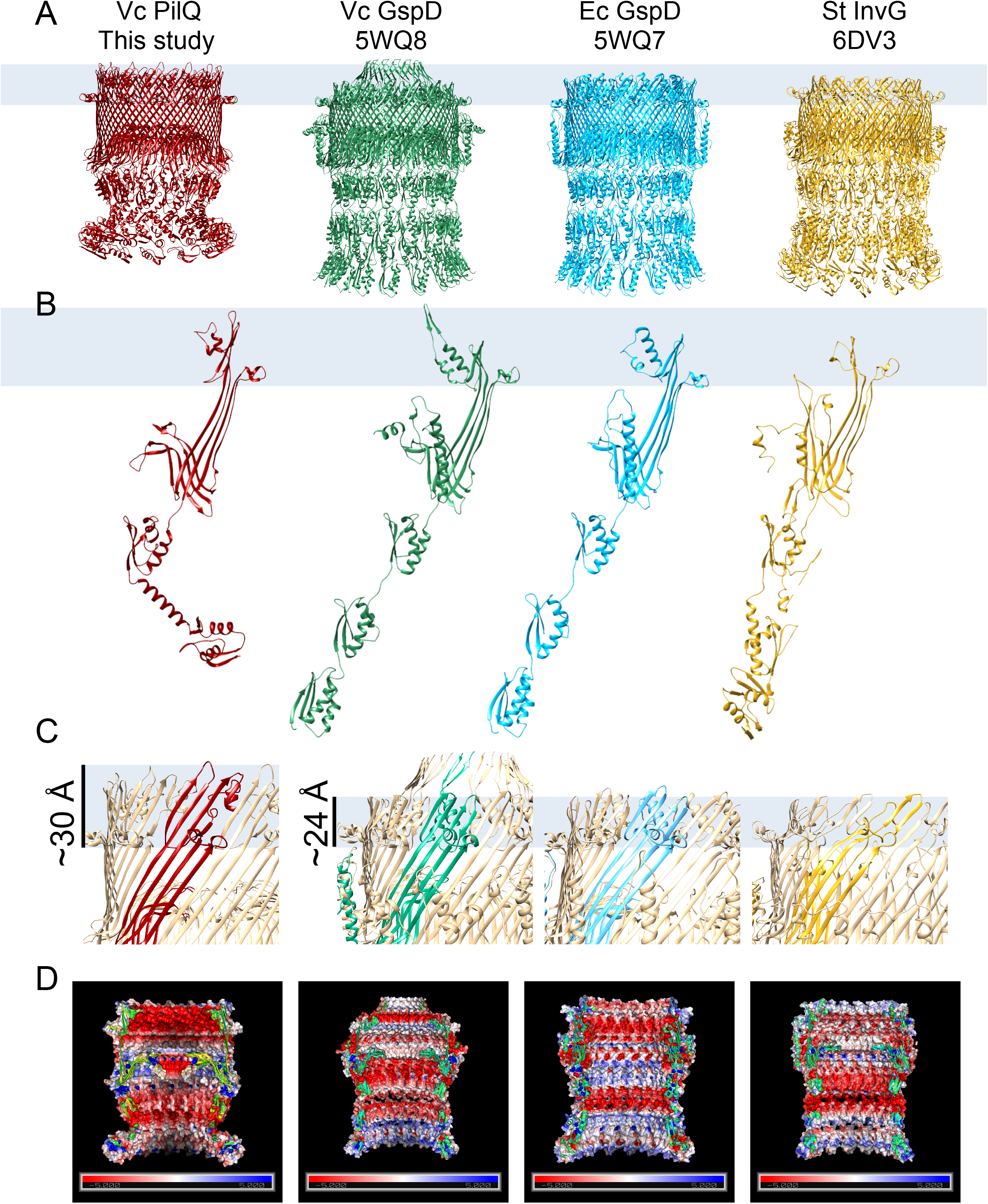
Comparison of VcPilQ to T2SS and T3SS secretins. The structures of VcPilQ (this study, dark red), *V. cholerae* GspD (5WQ8) (green), *E. coli* K12 GspD (5WQ7) (blue), and *S. typhimurium* InvG (6DV3) (yellow) (Jurrus et al. 2018; Worrall et al. 2016; Yan et al. 2017) are compared. The putative outer membrane location is depicted as a blue rectangle. The structures are shown as multimers **(A)** or monomers **(B).** In **(C)**, the multimer is shown in tan with one subunit colored. The outer membrane region is highlighted to show that VcPilQ (left, red) has a ∼30 Å membrane spanning distance, while VcGspD, EcGspD, and StInvG have about a 24 Å membrane spanning distance. **(D)** Adaptive Poisson-Boltzmann Solver was used to calculate the electrostatic potential calculation of each secretin (Jurrus et al. 2018). The inner cavity of each secretin is shown. The scale varies from −5 (red) to +5 (blue) in units of K_b_T/e_c_.

While their structures are similar, the electrostatic characteristics of the inner surfaces of the T4P, T2SS and T3SS secretins vary (**Figure 3D**). The inner beta-lip, N3, and coil regions of PilQ (**Figure 1C**) are all highly negatively charged (**Figure 3D**). In contrast, in the *V. cholerae* GspD and *E. coli* K12 GspD structures, there are alternating negatively- and weakly positively-charged regions. These differences are likely related to the function of PilQ in natural competence: the DNA cargo of the Type IV competence pilus is also negatively charged, so it is possible that this electrostatic repulsion will help the cargo pass through the cavity, rather than getting stuck. By comparison, for T2SS secretins, the charge alternates in the inner cavity.

### The N0 domain in VcPilQ agrees with previous structures

The N0 domain of VcPilQ (residues 160 to 227) was resolved 4 to 7 Å local resolution. This allowed us to build a model based on homology to previously solved structures of N0 domains (**Figure 1C**), which includes a crystal structure of isolated N0 domain from *Neisseria meningitidis* (4AR0) ^20^ (**Supplemental Figure 9**).

Only one of the seven T2SS secretins, and one of the T3SS published structures resolves the N0 domain (Worrall et al. 2016, Chernyatina and Low 2019). Here we can resolve the N0 domain (residues 160 to 227) of PilQ to 4 to 7 Å local resolution, which allowed us to build a model based on homology to previously solved structures of N0 domains (the cross-linked *Klebsiella oxytoca* PulD ^38^ and *S. typhimurium* InvG ^30^), plus a crystal structure of an isolated N0 domain from *Neisseria meningitidis* PilQ (4AR0)^20^ (**Supplemental Figure 10**). Our VcPilQ N0 structure (**Figure 1C**) falls within 1.2 Å RMSD of each of the previous structures (**Supplemental Figure 10E**).

### VcPilQ uses a novel helical coil to transition into the N3 domain

In VcPilQ, a 32 Å alpha-helix follows the N0 domain (**Figure 1C**). None of the T2SS or T3SS structures contain a helical coil to link N-terminal domains; instead their periplasmic protein domains are linked by unstructured loops (**Figure 3**). This difference was not anticipated, as homology models of VcPilQ based on other structures more closely match the previously published T2SS and T3SS structures (**Supplemental Figure 7**).

Following the end of this helix, the protein chain abruptly changes direction (∼104° angle) as the coil flows into the N3 domain (**Figure 1C**). This dramatically reduces the channel diameter, from 90 Å at the bottom of the N0 domain to 60 Å across the N3 domain (**Figure 4**). In the T2SS and T3SS structures, the diameter of the channel is relatively constant.

**Figure 4:**
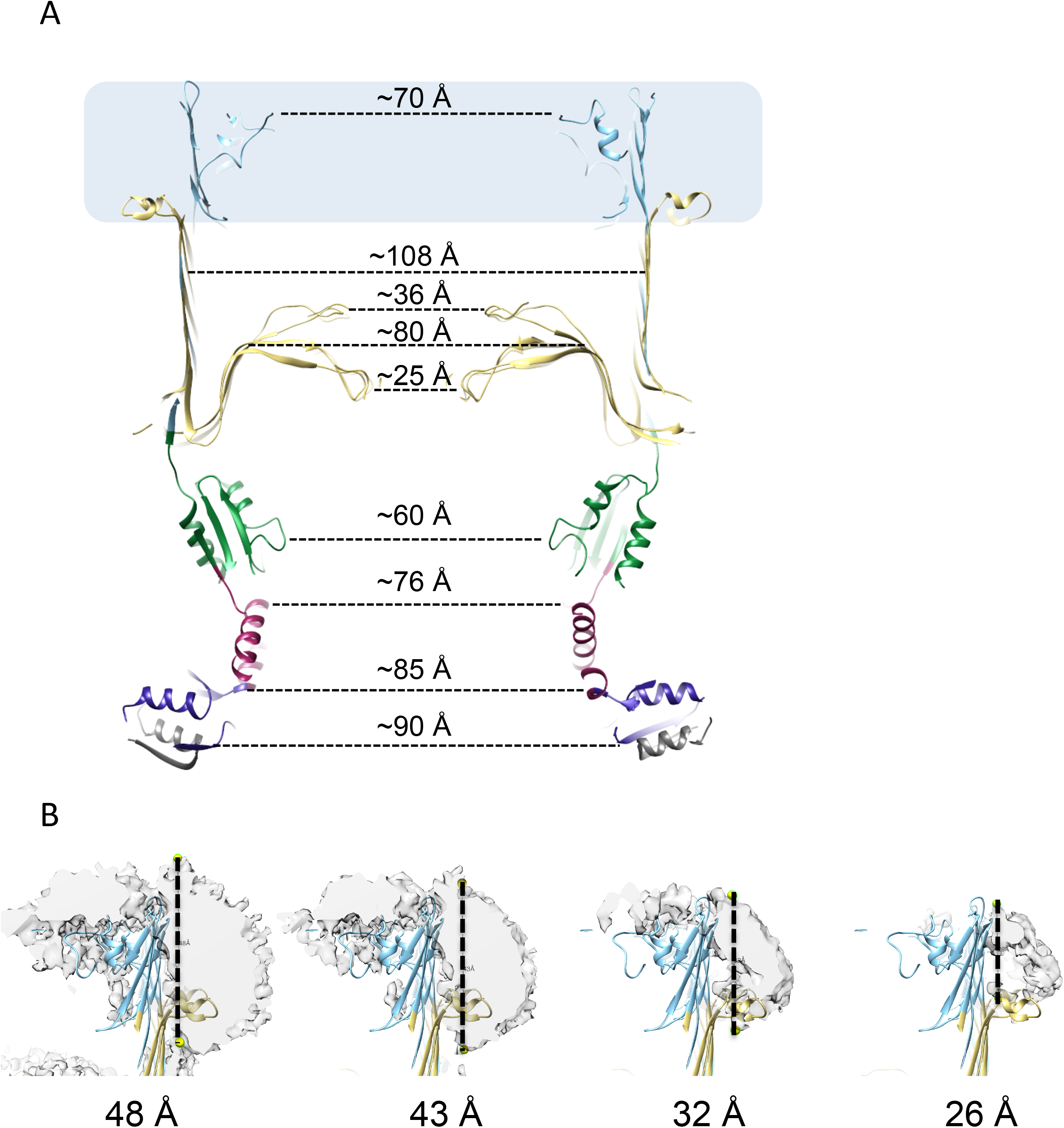
Dimensions of the VcPilQ inner channel and outer membrane region. **(A)** A central slice of the atomic model of *V. cholerae* PilQ colored as in Figure 1. The inner cavity distances are depicted with dashed lines and labeled in ångstroms. The outer membrane is depicted as a blue rectangle. **(B)** VcPilQ micelle density with putative outer membrane diameter labeled in Å. The non-protein cryoEM density of VcPilQ is shown in grey at four different Chimera thresholds (left to right). The atomic model of VcPilQ is shown in blue and yellow according to the color scheme of Figure 1. In each image, yellow spheres mark the approximate top and bottom of the micelle density. The distance between the yellow spheres (dashed line) is shown below each image in Ångstroms.

### The putative outer membrane region of VcPilQ is thicker than T2SS secretins

The secretin amphipathic helix lip (AHL) is thought to be a key determinant for secretin outer membrane insertion and among Type IV pilus secretins, the AHL is conserved ^18^. The AHL is thought to mark the lower boundary of the outer membrane region of secretins ^18, 29, 30^. The upper boundary of the outer membrane is less clear since the micelle density in cryoEM reconstructions does not always match the positions of aromatic residues ^25^.

In our VcPilQ atomic model, the distances between the bottom of the AHL to the top of the beta strands is about 3 nm. As seen in **Figure 3C**, this putative outer membrane region of VcPilQ is substantially taller than the same region in the previously published T2SS structures. To investigate more than just the residue locations, an inverted mask based on the atomic model density of VcPilQ was generated and subtracted from the empirical cryoEM density, which reflects unmodeled density in the cryoEM map that is not accounted for by the atomic model (**Figure 4C**). This presumed micelle density (in grey) blooms around the putative outer membrane region of the protein. This density appears both on the outside of the beta barrel, and inside, coating the inner lip of PilQ. It remains unclear which residues of PilQ are embedded within the outer membrane *in situ* (see Discussion).

### Cysteine mutants indicate gate must open for pilus biogenesis and natural transformation

PilQ is thought to mediate DNA uptake during natural transformation in *V. cholerae*. Blocking DNA uptake by inhibiting PilQ could prevent *V. cholerae* from undergoing horizontal gene transfer in chitin biofilms, which can promote the spread of antibiotic resistance genes and virulence factors. Toward this end, we designed cysteine pair mutants to reversibly lock the gate with disulfide bonds. The structure of VcPilQ was analyzed using the Disulfide by Design 2.0 web tool to identify residue pairs with geometries that could support a disulfide bond ^39–41^. The top hits from this screen were analyzed in Chimera and introduced into the *V. cholerae* genome. Two cysteine pairs (S448C/S453C and L445C/T493C) were able to assemble into PilQ multimers (**Figure 5**). The S448C/S453C pair is in the proximal hairpin of the gate, likely crosslinking adjacent PilQ monomers(**Figure 5C, 5E**). The L445C/T493C pair likely crosslinks the upper and lower gate hairpins of a single PilQ monomer(**Figure 5D, 5F**),. In both cases, under oxidizing conditions, the cysteine pair mutants demonstrated lower transformation efficiency than the control PilQ strain (**Figure 6A, Supplementary Figure 8**). Also, both mutants exhibited reduced piliation (**Figure 6B**). High levels of the reducing agent DTT were toxic to WT *V. cholerae* cells and inhibited piliation (**Figure 6B-D**) and natural transformation (**Supplementary Figure 8**). However, when the cells were grown with varying subinhibitory concentrations of DTT, the cysteine pair mutants recovered transformation efficiency and piliation similar to the control parent strain (**Figure 6C-D**). The S448C/S453C mutant is slightly more recalcitrant to DTT rescue, which may be due to the fact that these cysteines are located further down in the gate region making them less accessible to the reducing agent.

**Figure 5:**
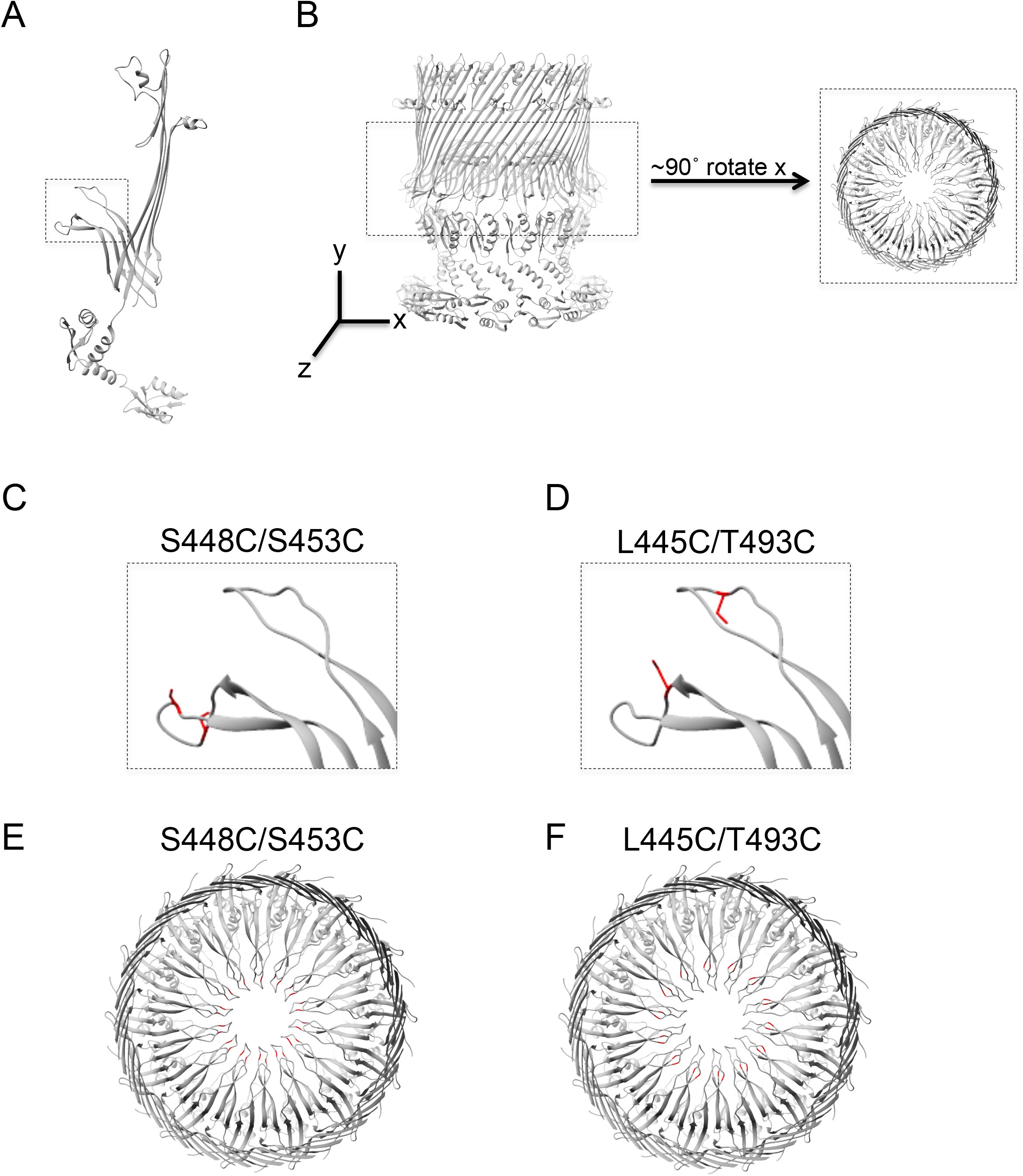
Design of cysteine pair mutants to lock the gate. The atomic model of VcPilQ is shown as one chain **(A)** or as a multimer **(B)**. In **(C)** and **(E)**, the S448C/S453C mutant, which locks the lower gate, is shown. In **(D)** and **(F)**, the L445C/T493C mutant, which links the upper and lower gates, is shown. In **(A)**, the upper and lower gate are highlighted in a dashed line box. A zoomed in region of this box is shown in **(C)** and **(D)** with the cysteine pair mutant residues highlighted in red. In **(B)**, the dashed line rectangle highlights the gate region, which is shown on the left after a 90° rotation about the x axis. The coordinate system is shown. This gate region is shown in (E) and (F) with the cysteine pair mutant residues highlighted in red.

**Figure 6:**
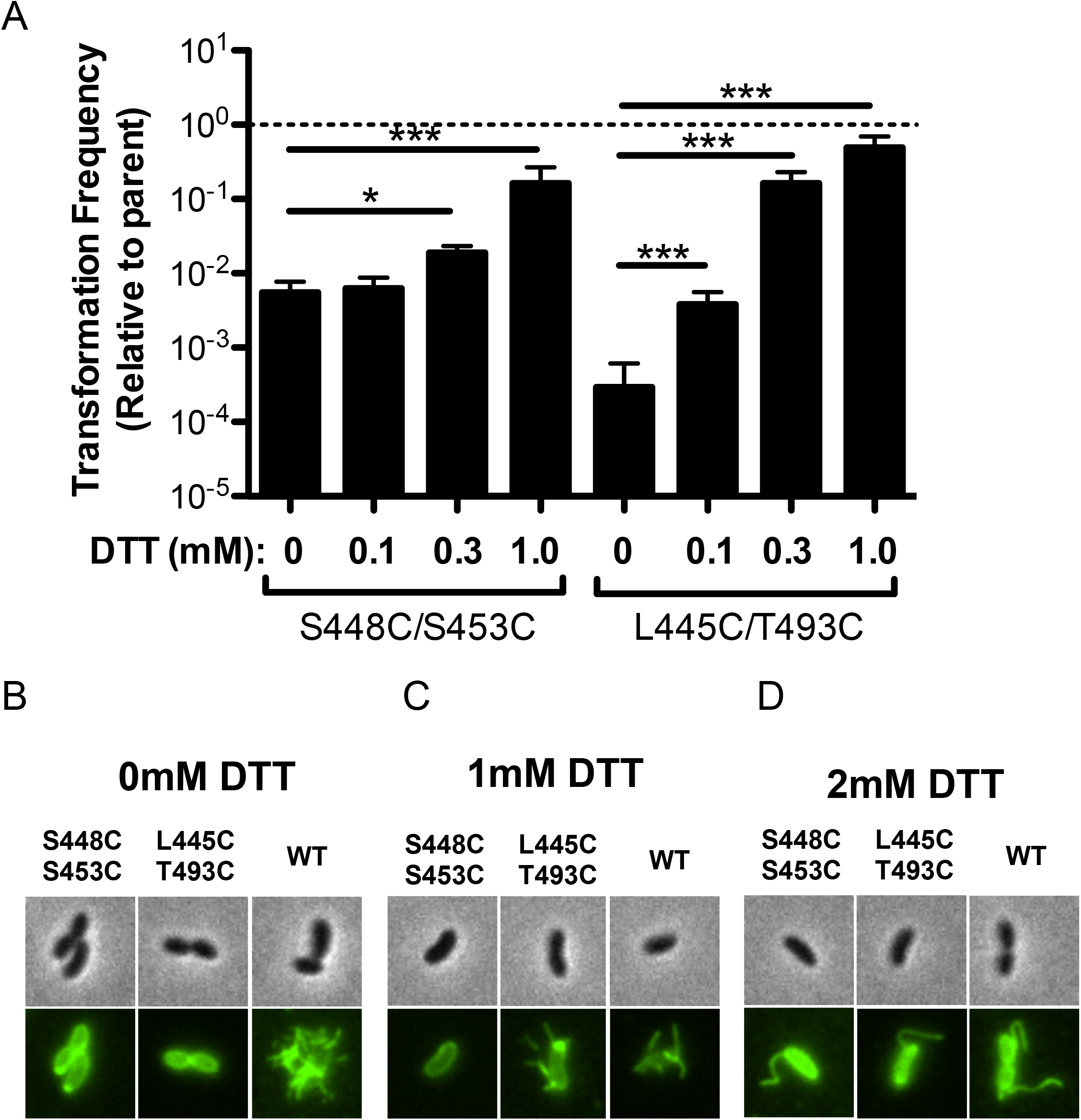
Cysteine pair mutants lock the gate, reduce piliation, and reversibly inhibit transformation. **(A)** The transformation frequency of two cysteine pair VcPilQ mutants (S448C/S453C on the left and L445C/T493C on the right) is plotted. Data are normalized to the parental strain that expresses wild-type VcPilQ. Natural transformation assays were performed in the presence of varying concentrations of DTT (0 to 1.0 mM). Data are from at least 4 independent biological replicates and shown as the mean ± SD. The dashed line indicates the transformation frequency expected if mutants are equivalent to the parental strain expressing wild-type VcPilQ. Statistical comparisons were made by one way ANOVA with Tukey’s post test. * = *p* < 0.05, *** = *p* < 0.001. **(B)-(D)** Representative phase contrast (top) and epifluorescence images (bottom) of *V. cholerae* pilA-Cys cells containing the indicated mutations in PilQ when grown in the presence of (B) 0mM, (C) 1mM, or (D) 2mM DTT prior to labeling with AF488-maleimide to visualize bacterial pili.

## Discussion

Here we present the first high-resolution structure of a bacterial Type IV competence pilus secretin. We observed key differences in the outer membrane region and the periplasmic region among the different members of the secretin family. These differences identify weaknesses in relying on homology models of evolutionarily related secretins, like the T2SS secretin GspD, to understand PilQ.

### Homology modeling was insufficient to predict a Type IV Pilus Secretin structure

Before we solved the VcPilQ structure, we used homology modeling to predict the structure of VcPilQ based on its sequence and the structures of previously solved secretins ^42^. The top five predictions are shown **Supplemental Figure 7**. Comparing these predictions to our now-solved structure reveal significant differences, including the thick outer membrane domain of VcPilQ and the extent of the alpha helix between the N0 and N3 domains (**Supplemental Figure 7**). Additionally, homology modeling cannot predict the three-dimensional arrangements and orientations of the protein domains, the number of subunits, or the inner barrel diameter. These features can be hypothesized based on T2SS or T3SS secretin structures, but that strategy would neglect the clear variations in selectivity and functionality across the Secretin superfamily. It is possible that structural variability between secretins also indicates which regions of the protein are more often repurposed by evolution to generate new function and/or selectivity.

### Outer membrane-spanning domain

Curiously, in some previous secretin structures, the trans-outer-membrane region appears to be only 2-3 nm thick (**Figure 3C**), which is much thinner than a typical membrane. This has left it unclear which regions of secretin molecules are actually embedded in the outer membrane, and whether any residues are exposed to the extracellular surface. To investigate these questions, we used the Positioning of Proteins in Membrane (PPM) web server to predict the location of the transmembrane region of VcPilQ ^43^. The PPM web server result delineates the hydrophobic region of the atomic model, which is shown by the dark horizontal lines parallel to the membrane in **Figure 7**. The full lipid bilayer membrane will then extend above and below these hydrophobic boundaries to account for the polar head groups of the lipids. As we predicted based on the micelle density (**Figure 4B**), the lower boundary of the hydrophobic region is the amphipathic helix lip. The upper hydrophobic boundary encompasses the beta strands of the beta lip, so the loops likely protrude into the polar head groups.

**Figure 7:**
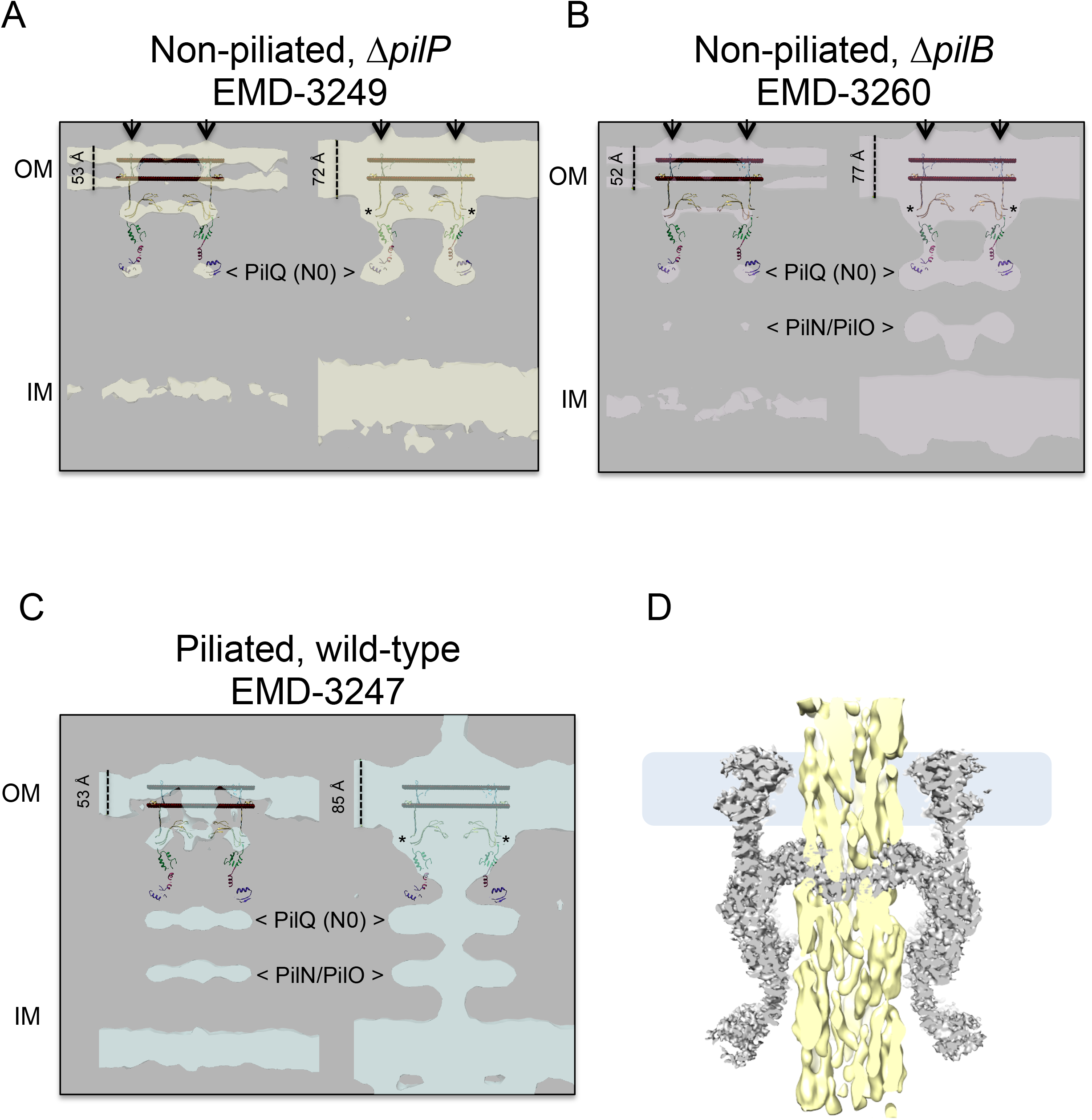
Comparison of VcPilQ with sub-tomogram averages of *M. xanthus* Type IVa pilus machinery, and with the *E. coli* Type IVa pilus. Sub-tomogram averages of the Type IVa pilus machinery in various *Myxococcus xanthus* strains are compared with VcPilQ^44^. VcPilQ is compared with the non-piliated strains Δ*pilP* (**A**, EMD-3249) and Δ*pilB* **(B**, EMD-3260), and with the piliated wild-type strain (**C,** EMD-3247). In the ΔpilP strain, only PilQ and TsaP are present in the complex. The putative TsaP location is marked with an asterisk in the rightmost volume in (**A), (B)** and **(C).** The location of the PilQ N0 domain is marked with arrowheads. In **(B)** and **(C)** the location of the MxPilN and MxPilO proteins are marked with arrowheads. Each Mx sub-tomogram average structure is depicted at a high cryoEM threshold (left) to delineate the inner and outer leaflets of the outer membrane, and at a lower cryoEM threshold (right) to fully reveal the protein features. For each cryoEM threshold, a dashed line is used to indicate the apparent membrane thickness, which ranges from 52 Å to 85 Å. The structure of the VcPilQ solved here is shown with coloring like in Figure 1. **(D)** A central slice of the cryoEM density of VcPilQ (grey) is shown with the *E. coli* Type IVa pilus structure (yellow) fit in ^52^. The outer membrane is depicted as a blue rectangle.

Next we docked the VcPilQ structure into a sub-tomogram average of a closely related PilQ imaged *in situ*. Recently, Chang et al. undertook an exhaustive cryoelectron tomography (cryoET) study of the T4aP machinery in *Myxococcus xanthus* (MxT4aP) ^44^. In the resulting sub-tomogram averages of the T4aP machinery, the location of each component protein was identified. The domain organization of MxPilQ is nearly identical to that of VcPilQ^45^ (**Supplemental Figure 11A**), but it has two more AMIN domains at its N-terminus (**Supplemental Figure 11A**). The Phyre2 protein model of MxPilQ also looks remarkably similar to the structure of VcPilQ solved here (**Supplemental Figure 11B)**^46^, even though the Phyre2 model was built independently.

We therefore docked the atomic model of VcPilQ into the sub-tomogram averages of the non-piliated MxT4aP in *M. xanthus* Δ*pilP* (EMD-3249), the non-piliated MxT4aP in *M. xanthus* Δ*pilB* (EMD-3260), and the piliated MxT4aP in wild-type *M. xanthus* (EMD-3247) (**Figure 7A-C**)^44^. Here we focus on the Δ*pilP* MxT4aP sub-tomogram average (EMD-3249) because in this mutant only PilQ and TsaP localize correctly, and are therefore the only proteins likely to be present in the sub-tomogram average (**Figure 7A**)^44^. The putative location of TsaP suggested in Chang *et al.* 2016 is marked with an asterisk in **Figure 7**. In *M. xanthus*, TsaP is a peptidoglycan-binding protein^47^. In *V. cholerae*, the TsaP homolog LysM has not been implicated in Type IV competence pilus function. The Δ*pilB* MxT4aP sub-tomogram average (EMD-3260), which lacks the extension ATPase *pilB*, is also shown because it was used to build a model of the non-piliated MxT4aP system (PDB 3JC9) in Chang *et al*. 2016 (**Figure 7B**).

The atomic model of VcPilQ was docked into the MxT4aP sub-tomogram averages using the predicted hydrophobic boundaries. At a high cryoEM density threshold, the lipid bilayer is clearly observed as two leaflets in the MxT4aP sub-tomogram averages (left in **Figure 7A-C**), so there is little doubt where the hydrophobic region of VcPilQ should be placed in the sub-tomogram average. The bilayer produces stronger features in the MxT4aP sub-tomogram averages than the protein, so a lower cryoEM density threshold is required to visualize the full MxT4aP machinery (right in **Figure 7**). In the non-piliated sub-tomogram averages (**Figure 7A-B**), VcPilQ agrees well with the MxPilQ density. The piliated MxT4aP clearly undergoes a conformational change to accommodate the pilus in which the distance between the inner and outer membranes increases (**Figure 7C**). In that case the VcPilQ N0 domain doesn’t extend far enough into the periplasm to line up with the MxPilQ N0 domain (labeled in **Figure 7C**).

The gate region of VcPilQ nicely superimposes onto the gate of the *M. xanthus* secretin, which suggests the VcPilQ structure was positioned correctly. However, the putative outer membrane region in VcPilQ does not extend across the entire outer membrane of the MxT4aP sub-tomogram averages. The thickness of the outer membrane in the sub-tomogram averages ranges from 52 to 85 Å depending on the cryoEM density threshold (labeled in **Figure 7A-C**), whereas the amphipol micelle region in VcPilQ seems to be between 25 and 50 Å thick (**Figure 4B**). While this discrepancy may be partially explained by the observation that the outer membrane is about 5 nm thick in *M. xanthus* and 4 nm thick in *V. cholerae*^48^, the shortness of the other predicted trans-membrane domains of secretins (**Figure 3C**) and the two bumps outside the outer membrane in the sub-tomogram average directly above the secretin barrel (arrows in **Figure 7A** and **B**) call into question whether and how parts of PilQ may penetrate the outer membrane *in situ*.

### VcPilQ could accommodate a Type IV Competence Pilus with gate rearrangement

The Type IV competence pilus machinery extends and retracts the Type IV competence pilus through the PilQ outer membrane pore. In our structure of VcPilQ in the absence of the pilus and accessory proteins, the inner diameter of the channel ranges from 25 to 108 Å (**Figure 4A**).

The *V. cholerae* Type IV competence pilus is a Type IVa pilus. The major pilin subunit in the T4aP is typically smaller than the major pilin subunits in the T4bP ^49^. CryoEM structures of several Type IVa Pili (T4aP) have been solved from *N. meningitidis* (NmT4aP, 6 Å resolution, 6 to 7 nm diameter), *N. gonorrhoeae* (NgT4aP, 5 Å resolution, 6 nm diameter), *E. coli* (EHEC strain, EcT4aP, 8 Å resolution, 6 nm diameter), and *P. aeruginosa* (PaT4aP, 8 Å resolution, 5 nm diameter) ^50–52^. Additionally, a recent preprint on bioRxiv studying Type IV pili in *T. thermophilus* demonstrated the presence of narrow (4.5 nm) and wide (7 nm) pili ^53^. In **Figure 7D**, the EcT4aP cryoEM structure (yellow) is fit into our VcPilQ structure (in grey)^52^. The gate region with diameters of 25 Å at the lower gate and 36 Å in the upper gate clashes with the pilus (**Figures 4A and 7D**). VcPilQ was solved in the absence of the pilus, so this gate conformation likely represents a “closed” state in cases where the pilus is absent or fully retracted. The next narrowest region in the VcPilQ channel is across the periplasmic N3 ring (∼60 Å in **Figure 4A**). Some of the previously solved T4aP are 6 to 7 nm in diameter, which would be a tight fit in this region. The N3 domain is connected to the secretin domain by a loop, so it could possibly expand to accommodate a 70 Å pilus.

We used homology modeling to predict the VcT4aP monomer structure, but because Neuhaus et al. reported both wide and narrow pili assemebled by pilins whose subunit structure are almost identical, we do not feel confident guessing the VcT4aP diameter^53^. Regardless, the gates in VcPilQ would have to move to accommodate a pilus. Consistent with this notion, our disulfide locked cysteine pair mutants could not extend pili in the absence of reducing agent, which strongly suggests that the conformation adopted in the presented structure (in which these cysteine pairs would be in close enough proximity to disulfide bond) represents the closed gate conformation and cannot accommodate a pilus fiber. Additionally, the sub-tomogram averaging of the MxT4aP demonstrates a secretin conformational change with and without a pilus present (**Figure 7B-C**)^44^.

In Yan *et al.*, a glycine in the VcGspD gate (G453) was identified as a putative hinge point to facilitate gate opening, and showed that the G453A mutant trapped the T2SS secretin in a partially open state^24^. In our VcPilQ structure, the inner channel distance between the corresponding glycine (G439) alpha carbons is about 8 nm (**Figure 4A**). Thus, we hypothesize that a gate hinge mechanism could accommodate pili up to 7 nm in diameter. In an open state the gate loops could flip up toward the extracellular space to accommodate the pilus.

## Conclusion

Here we report the first high-resolution structure of a Type IV competence pilus secretin, *V. cholerae* PilQ. This protein complex facilitates DNA uptake into diverse bacterial species to aid in their evolution, and thus, represents a potential target for therapeutic intervention. The *V. cholerae* Type IV competence pilus is a model system to study natural transformation in bacteria. We identify key differences between VcPilQ and the previously published structures of T2SS and T3SS secretins. We designed cysteine pair mutants to reversibly seal the VcPilQ gate and inhibit natural transformation, which can be used as a tool to further investigate the function of the Type IV competence pilus machinery *in situ*. We suggest a structural rearrangement that would transition our closed VcPilQ into a piliated state that could accommodate previously solved structures of Type IV pili. We also compare our structure to previous T4aP sub-tomogram averaging results in *M. xanthus* which call in question which parts if any of PilQ extend across the outer membrane. Together, these results elucidate the structure of PilQ and provide mechanistic insight into natural competence in *V. cholerae*.

---

**Supplemental Figure 1: Expression and purification of VcPilQ from *V. cholerae* cells**

**(A)** The functionality of N-terminally deca-histidine tagged native PilQ was assessed via natural transformation assays. These data indicate that the tagged variant supports equal rates of transformation compared to an isogenic parent strain, thus indicating that it is fully functional. Data are from four independent biological replicates and shown as the mean ± SD. Statistical comparisons were made by unpaired Student’s t-test. NS = not significant. **(B-F)** Gel electrophoresis analysis of VcPilQ in amphipols by coomassie stain **(B)**, **(D), (E)**, and western blotting against anti-His tag **(C)**, **(F)**. The multimer band is indicated with an asterisk.

**Supplemental Figure 2: *V. cholerae* PilQ sequence with mass spectroscopy results.**

Mass spectrometry coverage of purified PilQ: The sequence of VcPilQ (571 amino acids) is shown in black with residue numbers marked ever 10 residues. Regions of the sequence corresponding to protein motifs (Signal Peptide, AMIN Domain, STN, Coil, Secretin_N and Secretin) or folded regions of the protein structure (N0, N3) are highlighted with flags below the relevant sequence. The sequence coverage achieved by mass spectrometry is highlighted with grey flags labeled “MS”. Approximately 65% of the sequence was represented in the peptides analyzed by mass spectrometry. The protein sequence was visualized using the Benchling Biology Software.

**Supplemental Figure 3: CryoEM data**

**(A)** Representative micrograph with a 10 Å low pass filter applied. A representative Fourier transform and contrast transfer function (CTF) fit of the micrograph is shown in the upper left corner. Scale bar, 200 Å. **(B)** Representative 2D classes calculated in Relion. The 2D class box is 441.6 Å wide. **(C)** A top view 2D class from demonstrates that VcPilQ has C14 symmetry. Yellow lines point to inner spokes in the PilQ gate. Each monomer is numbered in white.

**Supplemental Figure 4: CryoEM data processing flow chart**

The major steps in data processing are summarized in a flow chart.

**Supplemental Figure 5: Fourier Shell Correlation and 3DFSC of VcPilQ**

**(A)** Fourier shell correlation (FSC) of VcPilQ cryoEM structure. **(B) to (D)** The Salk institute 3DFSC server was used to examine the resolution in the X, Y and Z dimensions in the VcPilQ structure(Tan et al. 2017). The sphericity is 0.962 out of 1.

**Supplemental Figure 6: Local resolution of VcPilQ**

ResMap was used to calculate the local resolution per voxel of the density map. Resolution is plotted from 2.4 Å (cyan) to 8.4 Å (black). VcPilQ is shown from the side **(A),** as a central slice **(B),** from the top (extracellular side, **C**) and from the bottom (periplasmic side, **D**).

**Supplemental Figure 7: Homology modeling of VcPilQ**

I-TASSER was used to predict the structure of *V. cholerae* PilQ. I-TASSER reported five potential models (labeled VcPilQ Homology model 1 to 5 and shown with gray background). The ITASSER models are based on structures deposited in the PDB, so the five models can be sorted by their similarity to the T2SS with the cap (Vibrio-type), without the cap (Klebsiella-type) or the T3SS, so the models are shown next to the corresponding secretin type. On the right the structure of VcPilQ solved in this paper is shown in dark red. The membrane is blue.

**Supplemental Figure 8: Raw transformation frequency data for cysteine pair mutants.**

Transformation assays were performed with the indicated strains in the presence of DTT as indicated above the bars. Data are from at least 4 independent biological replicates and shown as the mean ± SD.

**Supplemental Figure 9: RMSD of VcPilQ versus published secretin structures.**

The monomer structure of VcPilQ (dark red) was aligned to the *E. coli* K12 GspD (5WQ7) **(A)**, the *V. cholerae* GspD (5WQ8) **(B)**, and the *S. typhimurium* InvG (6DV3 and 5TCQ) **(C-D)** (Worrall et al. 2016; Yan et al. 2017; J. Hu et al. 2018). Only the N3 and Secretin domains were compared. RMSD was calculated with MatchMaker in Chimera. The RMSD results are summarized in Table (**E).**

**Supplemental Figure 10: Alignment of VcPilQ N0 domain compared to others.**

The VcPilQ coil and N0 domain (red, residues 160 to 228) were compared to previously published N0 structures. The *N. meningitidis* PilQ N0 NMR structure (purple, residues 350-417) is shown fit to the VcPilQ N0 (red) with the smallest RMSD structure (0.9 Å) in **(A**) and with all structures (RMSD 0.9 to 1.2 Å) in (**B).** (**C)** *S. typhimurium* InvG (PDB 6DV3, residues 34-104), RMSD 1.1 Å. (**D)** The N0 domain of *Klebsiella oxytoca* PulD (PDB 6HCG, residues 27-100) RMSD 1.1 Å (Berry et al. 2012; Chernyatina and Low 2019; Worrall et al. 2016). Scale, 10 Å. (**E)** Table of RMSD results comparing VcPilQ N0 domain to other N0 domains.

**Supplemental Figure 11: Comparison of VcPilQ with putative *M. xanthus* PilQ structure**

**(A)** The domain organization of *V. cholerae* PilQ (top) and *M. xanthus* PilQ (bottom) calculated in CDVist (Adebali, Ortega, and Zhulin 2015). **(B)** The structure of VcPilQ (this study) in dark red is compared to the Phyre2 homology modeling prediction of the MxPilQ structure (right, pink)(Kelley et al. 2015).

**Supplemental Movie 1: Atomic model of VcPilQ fit into the cryoEM density**

A single subunit of VcPilQ (labeled and colored by domain as in Figure 1) is shown first by itself, then in the multimer. Next the model is displayed in the cryoEM density to demonstrate agreement.

## Methods

### Bacterial Strains and culture conditions

All *V. cholerae* strains were derived from the El Tor strain E7946^54^. Strains were routinely grown in LB Miller broth and agar. VcPilQ was purified from *V. cholerae* with a deca-histidine tag added to the N-terminus of PilQ at the native locus. The full genotype of this strain (TND1751) is 10xHis-PilQ, ΔVC1807::SpecR, lacZ::lacIq, comEA-mCherry, ΔluxO, Ptac-tfoX, ΔTCP::ZeoR, ΔMSHA::CarbR, ΔCTX::KanR.

For testing the impact of cysteine pair mutants on natural transformation, *V. cholerae* strains were generated where the native copy of PilQ was deleted and the corresponding PilQ allele was expressed at an ectopic site. The full genotype of the parent strain (TND2140) was ΔlacZ::Pbad-10XHis-PilQ CmR, ΔpilQ::TetR, ΔCTX::KanR, ΔMSHA::CarbR, ΔluxO, ΔTCP::ZeoR, comEA-mCherry, Ptac-tfoX. The cysteine pair mutants were isogenic other than the cysteine mutations introduced into the Pbad-10XHis-PilQ construct, which were PilQ S448C S453C (TND2169) and PilQ L445C T493C (TND2170).

To test the impact of cysteine pair mutants on pilus biogenesis, *V. cholerae* strains were generated akin to those described above, with the exception that the retraction ATPase PilT was deleted and the strains contained a cysteine substitution mutation in the major pilin that allows for competence pilus labeling as previously described^15^. The full genotype of the parent strain (TND2244) was ΔlacZ::Pbad-10XHis-PilQ CmR, ΔpilT::TmR, ΔpilQ::TetR, ΔCTX::KanR, ΔMSHA::CarbR, ΔluxO, ΔTCP::ZeoR, pilA S67C, comEA-mCherry, Ptac-tfoX. The cysteine pair mutants were isogenic other than the cysteine mutations introduced into the Pbad-10XHis-PilQ construct, which were PilQ S448C S453C (TND2242) and PilQ L445C T493C (TND2243).

All strains were generated by natural transformation and cotransformation exactly as previously described ^55^.

### Natural transformation assays

Natural transformation assays were performed exactly as described in ^15^. For reactions where strains harbored Pbad-10XHis-PilQ constructs, arabinose was added to a final concentrations of 0.2%. Where indicated, DTT was added at the indicated concentrations throughout the assay.

### Competence pilus labeling and microscopy

Cells were labeled with AlexaFluor 488-maleimide dye and imaged to visualize competence pili exactly as previously described ^15, 56^. All strains were grown with arabinose added to a final concentrations of 0.2% to induce expression of the Pbad-10XHis-PilQ construct. Where indicated, cells were grown in the presence of the indicated concentration of DTT prior to labeling.

### Expression

*Vibrio cholerae* expressing His-tagged PilQ (10xHis-PilQ, ΔVC1807::SpecR, lacZ::lacIq, comEA-mCherry, ΔluxO, Ptac-tfoX, ΔTCP::ZeoR, ΔMSHA::CarbR, ΔCTX::KanR) was streaked on Luria Broth agar plates and grown overnight at 30°C. Small cultures were seeded (5 mL) and grown over night at 30°C. The next day, 500 mL cultures were seeded with the 5 mL culture. LB broth was supplemented with 20 mM MgCl_2_, 10 mM CaCl_2_ and 100 µM IPTG. The large cultures were grown overnight at 30°C in beveled flasks. The following day, cultures were spun down (6,000 rpm, 4°C, 20 minutes) and the cell paste was weighed, aliquoted, and stored at −80°C.

### Purification

Cell pellet (15 g) was resuspended in lysis buffer (50 mM Tris HCl, pH 8, 300 mM NaCl, 1% DDM, 20 mM imidazole) supplemented with lysozyme (40 mg/mL in 50% glycerol/water), DNAse I (4 mg/mL in 50% glycerol/water), and EDTA-free Protease Inhibitor tablet (Roche, 11697498001). Lysis proceeded with stirring at 4°C for 20 hours. Lysate was clarified by ultracentrifugation (Beckman L8-M ultracentrifuge, Rotor Type 45 Ti, 35,000rpm, 1 hour). The supernatant was mixed with Ni NTA agarose beads (Anatrace, SUPER-NINTA25) and incubated with stirring (4°C, 8 hours). In a gravity column at 4°C, proteins conjugated to Ni NTA agarose beads were washed (50 mM Tris HCl, pH 8, 300 mM NaCl, 0.05% DDM, 70 mM imidazole), (50 mM Tris HCl, pH 8, 300 mM NaCl, 0.05% DDM, 300 mM imidazole), and eluted (50 mM Tris HCl, pH 8, 300 mM NaCl, 0.05% DDM, 1 M imidazole). Eluant was concentrated to ∼1 mg/L (EMD Millipore Amicon Ultra-15, 30 kDa cutoff, UFC903024). Concentrated PilQ (150 µL of ∼1 mg/mL protein) was exchanged into Amphipol A8-35 (0.585 mg for a 3:1 ratio, Anatrace, A835) and allowed to incubate at 4 C for 1 hr. Excess DDM was removed using Bio-Beads SM2 (Bio-Rad, 1523920) by incubating overnight at 4°C. The protein was concentrated.

### Electron microscopy

For cryoEM, Quantifoil R2/2 300 Mesh grids (EMS, Q33100CR2) were glow discharged (Pelco EasiGlow, 20 mA, 60 seconds). PilQ in amphipol (3 µL of ∼0.8 mg/L) was frozen on a Mark IV Vitrobot (FEI, 20 °C, 100% relative humidity, blot force −6, blot time 4 s). Micrographs were collected on a 300 kV Titan Krios microscope (FEI) with energy filter (Gatan) and equipped with a K3 direct electron detector (Gatan). Data was collected using Serial EM software with a pixel size of 1.104 Å (81,000x magnification) and a defocus range from −1.0 µm to −3.0 µm ^57^. A fluence of 19.8 electrons/pixel/second was used with a 3.7 s exposure time to collect 60 e-/ Å^2^.

### Image Processing

The cryoEM image processing workflow is summarized in **Supplemental Figure 4**. MotionCor2 was used for motion correction and dose weighting of 3,808 movies^58^. CTF correction was used to evaluate micrograph quality(Rohou and Grigorieff 2015). CryoSPARC blob picking on 2,510 micrographs yielded 3,100,353 potential particles^60^. After inspection, the 252,319 particles were analyzed by several rounds of 2D classification and 3D classification to yield 100,543 particles. These particles were moved to Relion using the UCSF PyEM package (https://github.com/asarnow/pyem) script^61^. In Relion, several rounds of 3D refinement, polishing and CTF refinement were used^62–64^. ResMap was used to calculate local resolution^35^.

### Model Building and Refinement

The initial model (residues 230-571) was auto-built using Buccaneer^65^. Subsequent building and model adjustments were performed by hand using COOT^66^. A homology model of the N0 domain (residues 160-229) was created using I-TASSER and manually docked using COOT^42, 67, 68^. Coulombic potential density for residues 1-159 were not observed. The model was refined in PHENIX version 1.16-dev3549 using phenix.real_space_refine with the resolution set to 3 Å ^69^. NCS constraints were applied for the 14 subunits and were automatically detected and refined. Automatically determined secondary structure restraints, rotamer restraints, and Ramachandran restraints were applied as well. The quality of the model was evaluated using EMRinger^70^ and Molprobity ^71^ (Table 1).

**Table 1.** Summary of single-particle data collection, 3D reconstruction, and model refinement

|  |  |
| --- | --- |
| **Imaging parameters and 3D reconstruction** |  |
| Acceleration voltage (kV) | 300 |
| Magnification (X) | 81,000 |
| Pixel size (Å) | 1.104 |
| Frame rate (s<sup>-1</sup>) | 0.092 |
| Exposure time (s) | 3.7 |
| Total exposure (e<sup>-</sup> / Å) | 60 |
| Particles |  |
| Micrographs used for selection | 2,510 |
| Defocus range (µm) | -0.5 to -3.5 |
| Windowed | 252,319 |
| In final 3D reconstruction | 100,543 |
| Resolution |  |
| ‘Gold-standard’ at FSC 0.5 (Å) | 3.0 Å |
| ‘Gold-standard’ at FSC 0.143 (Å) | 2.7 Å |
| Map-sharpening B factor (Å<sup>2</sup>) | -69 |
| **Model refinement** |  |
| Resolution in phenix.real space refine (Å) | 3.0 |
| Model-to-map fit (CC_mask) | 0.745 |
| Number of atoms/residues/molecules |  |
| NCS restrained chains | 14 |
| Protein atoms, residues (per chain) | 43736, 412 |
| Ramachandran angles (%) |  |
| Favored | 92.21 |
| Allowed | 7.79 |
| Outliers | 0 |
| r.m.s. deviations |  |
| Bond lengths (Å) | 0.006 |
| Bond angles (°) | 0.847 |
| Molprobity |  |
| Score | 2.81 |
| Clashscore | 10.53 |
| Rotamer outliers (%) | 10.91 |
| EMRinger score | 3.13 |

### Mass Spectrometry

After running a BioRad Stain Free gel and performing a coomassie staining, the band of interest was excised with a clean razor blade. The gel piece was destained with ammonium bicarbonate and reduced with DTT (50°C, 30 minutes). Next the sample was alkylated with iodoacetamide (room temperature, dark, 20 minutes).

The gel pieces were then dehydrated. Trypsin was used to digest the protein in the gel (37°C, overnight). Peptides were extracted from the gel matrix, dried, and de salted with a zip tip.

The in-gel-digested samples were subjected to LC-MS/MS analysis on a nanoflow LC system, EASY-nLC 1200, (Thermo Fisher Scientific) coupled to a QExactive HF Orbitrap mass spectrometer (Thermo Fisher Scientific, Bremen, Germany) equipped with a Nanospray Flex ion source.

Samples were directly loaded onto a C18 Aurora series column (Ion Opticks, Parkville, Australia). The 25cm x 50µm ID column (1.6 µm) was heated to 45° C. The peptides were separated with a 60 min gradient at a flow rate of 350 nL/min. The gradient was as follows: 2–6% Solvent B (3.5 min), 6-25% B (42.5 min), and 25-40% B (14.5min), to 100% B (1min) and 100% B (12min). Solvent A consisted of 97.8% H2O, 2% ACN, and 0.2% formic acid and solvent B consisted of 19.8% H2O, 80% ACN, and 0.2% formic acid.

The QExactive HF Orbitrap was operated in data dependent mode. Spray voltage was set to 1.8 kV, S-lens RF level at 50, and heated capillary at 275 °C. Full scan resolution was set to 60,000 at m/z 200. Full scan target was 3 × 10^6^ with a maximum injection time of 15 ms (profile mode). Mass range was set to 300−1650 m/z. For data dependent MS2 scans the loop count was 12, target value was set at 1 × 10^5^, and intensity threshold was kept at 1 × 10^5^. Isolation width was set at 1.2 m/z and a fixed first mass of 100 was used. Normalized collision energy was set at 28. Peptide match was set to off, and isotope exclusion was on. Ms2 data was collected in centroid mode.

Raw data were analyzed using MaxQuant (v. 1.6.5.0)^72, 73^. Spectra were searched against UniProt *V. cholerae* entries (3784 sequences) and a contaminant protein database (246 sequences). Trypsin was specified as the digestion enzyme and up to two missed cleavages were allowed. Precursor mass tolerance was 4.5 ppm after recalibration and fragment mass tolerance was 20 ppm. Variable modifications included oxidation of methionine and protein N-terminal acetylation. Carbamidomethylation of cysteine was specified as a fixed modification. A decoy database was used to set score thresholds to ensure a 1% false discovery rate at the protein and peptide level. Protein abundances were estimated using iBAQ and the fractional abundance was calculated as the protein abundance divided by the sum of all non-contaminant protein abundances^74^.

## Author Contributions

S.J.W. conceptualized the project, expressed and purified the protein, prepared samples for cryoEM, collected cryoEM data, processed cryoEM data, assisted in atomic model building, interpreted results, designed figures, and wrote the paper.

M.S. purified protein, assisted with cryoEM sample prep and data collection, built the atomic model, interpreted results, and provided feedback on the paper.

T.D. engineered the *V. cholerae* constructs, performed microbial assays, and interpreted results.

A.D. conceptualized the project, obtained funding, engineered the *V. cholerae* constructs, performed microbial assays, interpreted results, designed figures, and provided feedback on the paper.

G.J.J. conceptualized the project, obtained funding, and provided feedback on the paper.

## Acknowledgements

Cryo Electron microscopy was performed in the Beckman Institute Resource Center for Transmission Electron Microscopy at Caltech. Dr. Songye Chen and Dr. Andrey Malyutin assisted with data collection. Dr. Spiros D. Garbis, Dr. Annie Moradian, Dr. Michael Sweredoski, and Dr. Brett Lomenick at the Caltech Proteome Exploration Laboratory (PEL) performed and analyzed mass spectrometry results. Dr. Naima Sharaf, Jeffery Lai, and Prof. Doug Rees provided invaluable advice on membrane protein biochemistry and instrumentation. Jane Ding and Welison Floriano provided computational support. Dr. Debnath Ghosal, Dr. Mohammed Kaplan, Dr. Davi Ortega, Dr. Catherine Oikonomou, Dr. Lauren Ann Metskas, Dr. Christopher Barnes, Claudia Jette, and Andrew Schacht provided feedback and advice. This work was supported in part by grant R35GM128674 from the National Institutes of Health to A.B.D.

