## Supplemental Figures 1 to 11 for "CryoEM structure of the Vibrio cholerae Type IV competence pilus secretin PilQ"

2020

Supplemental Figures 1-11

### Supplemental Figure 1

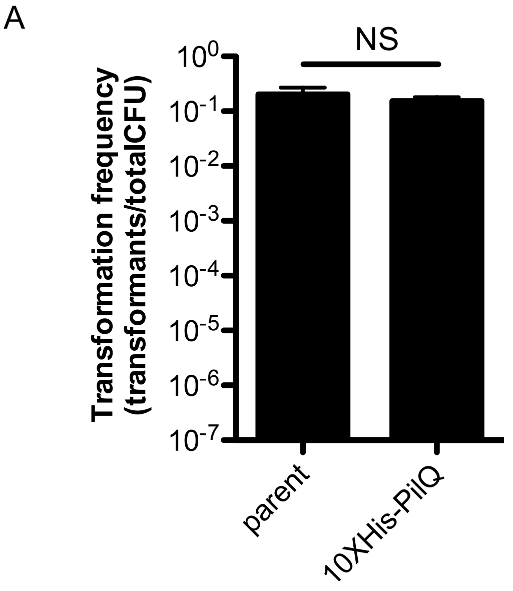


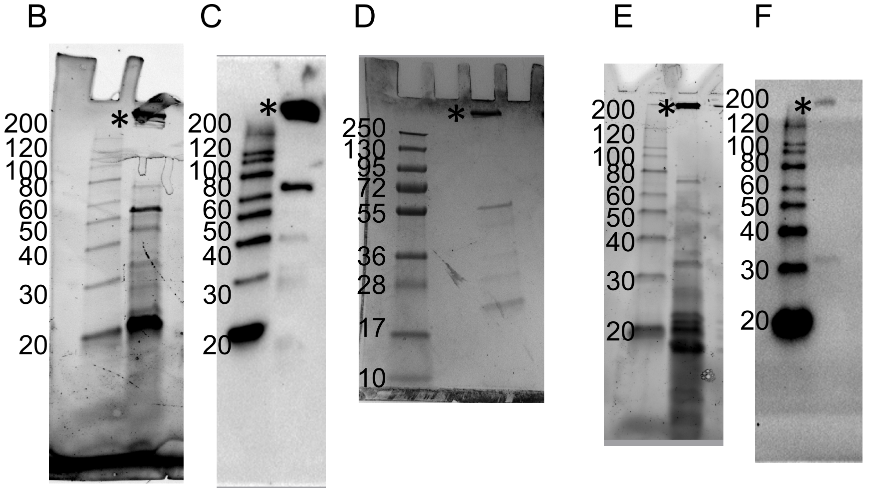

### Supplemental Figure 2

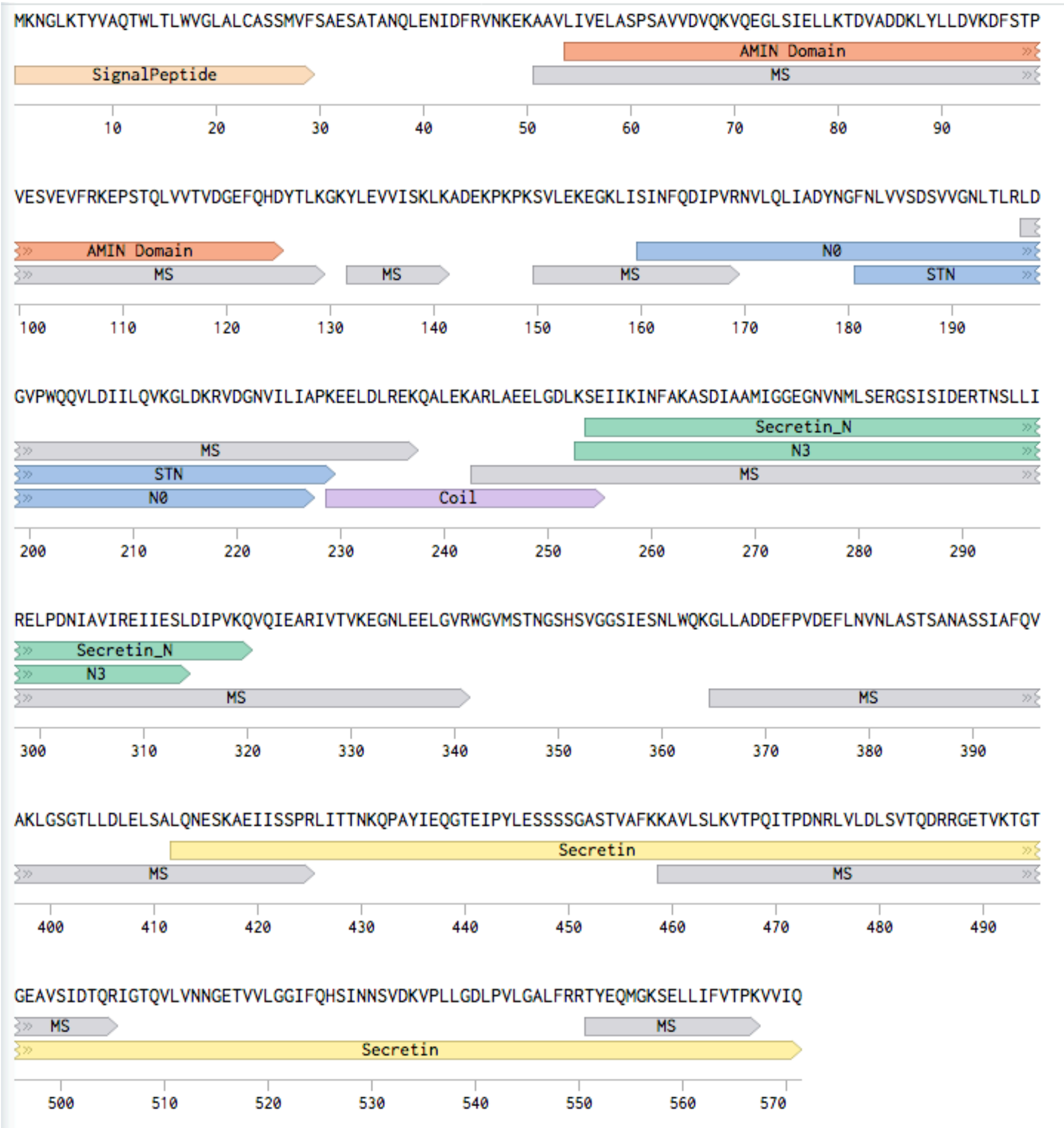

### Supplemental Figure 3

A

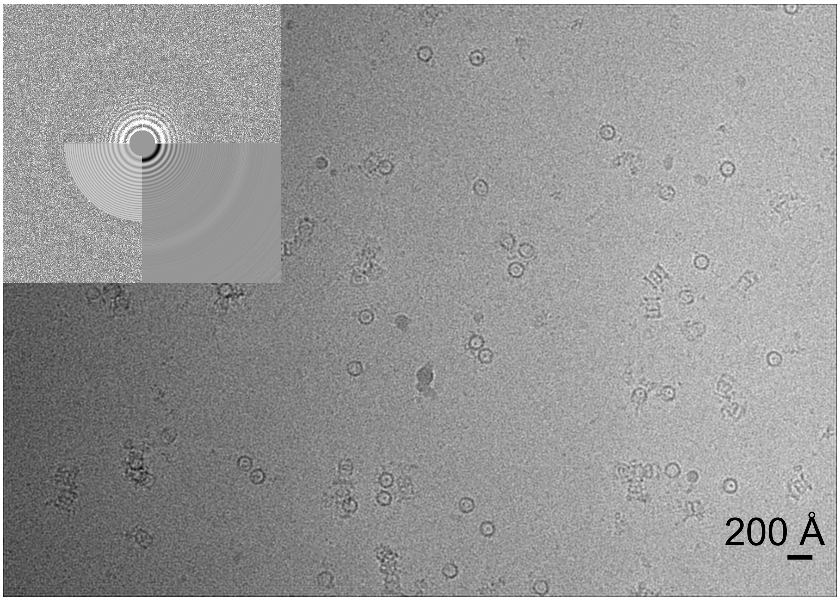


B

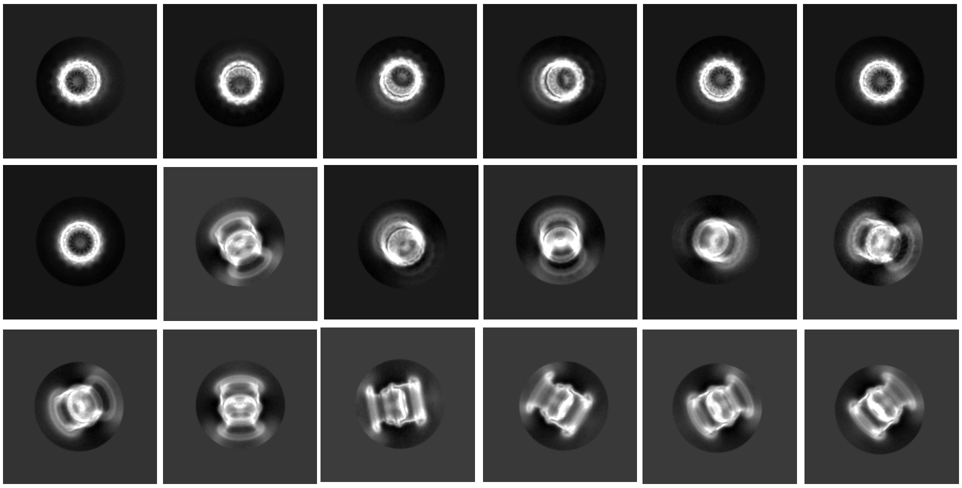

C

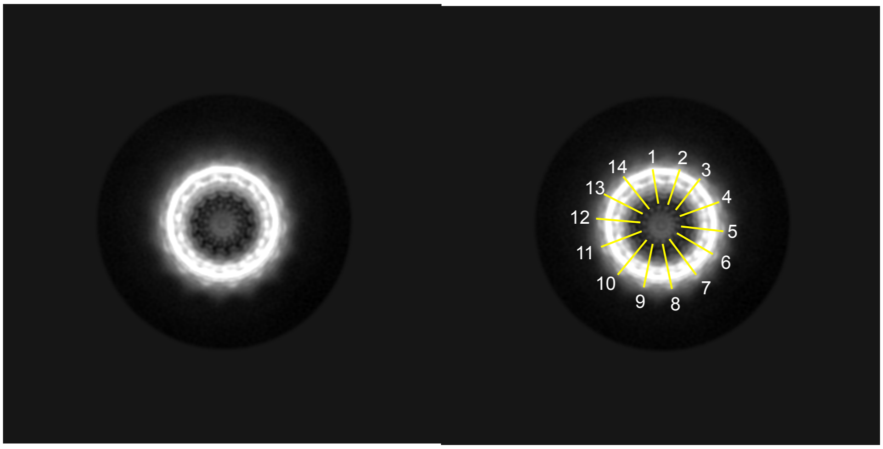

### Supplemental Figure 4

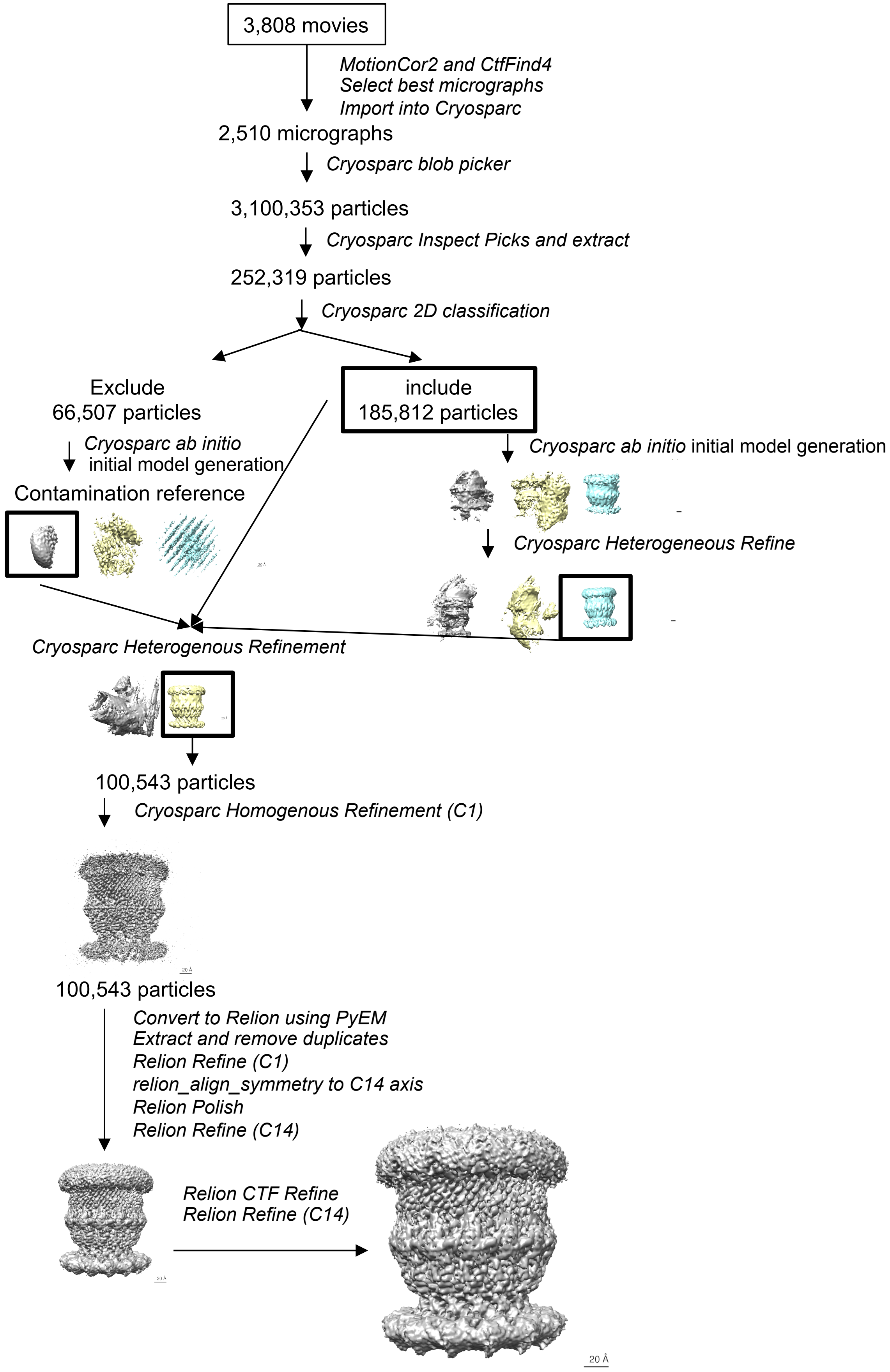

### Supplemental Figure 5

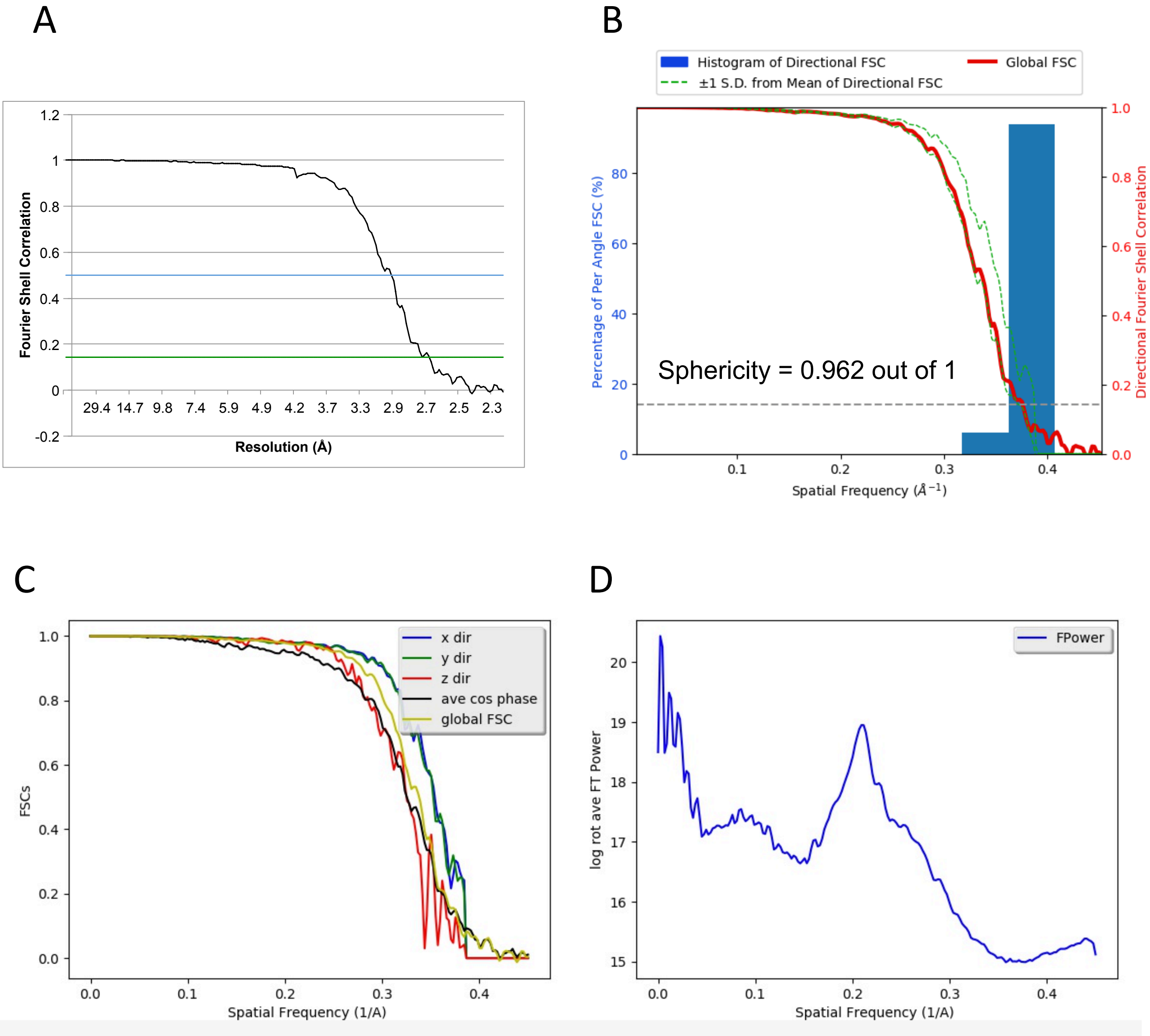

Supplemental Figure 6

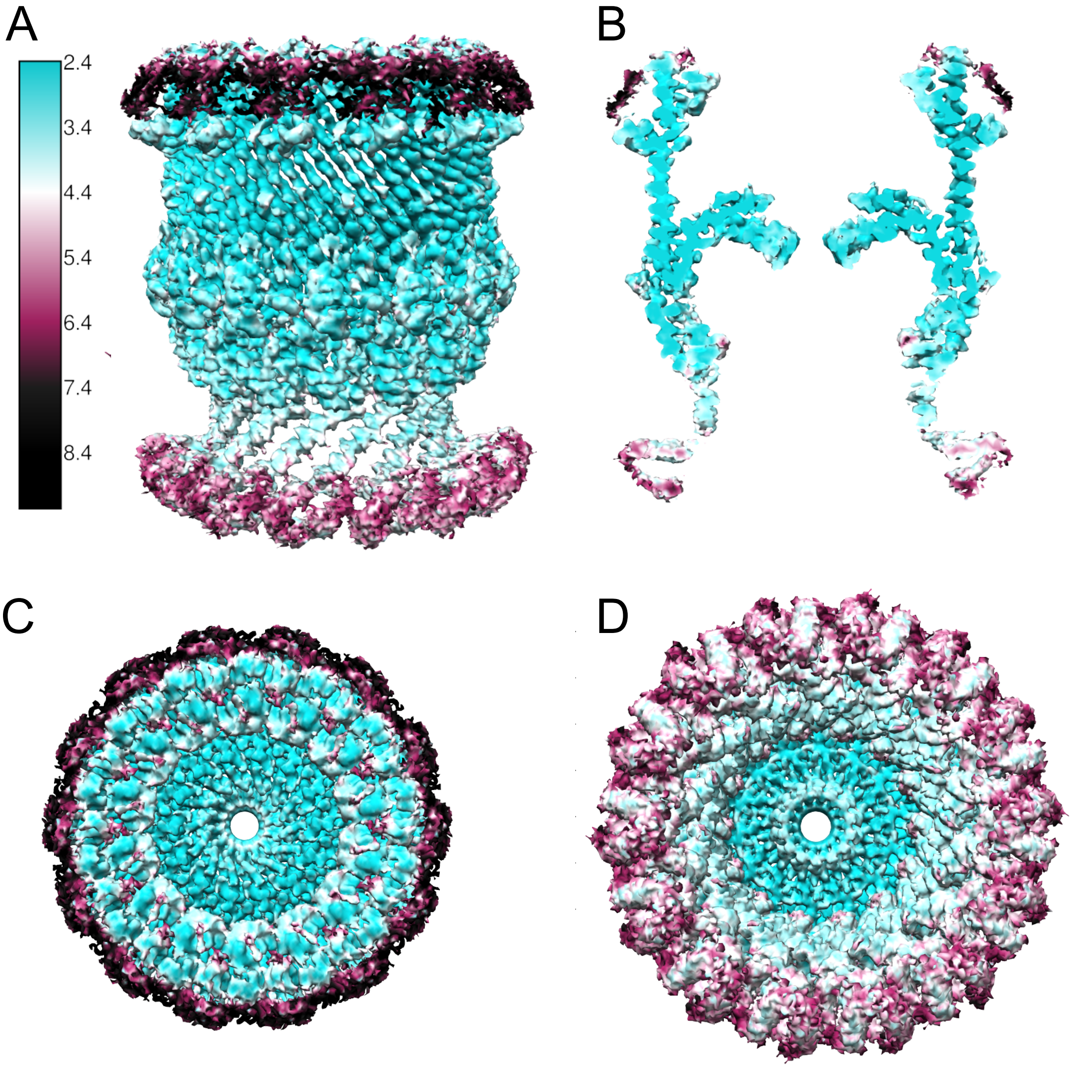

### Supplemental Figure 7

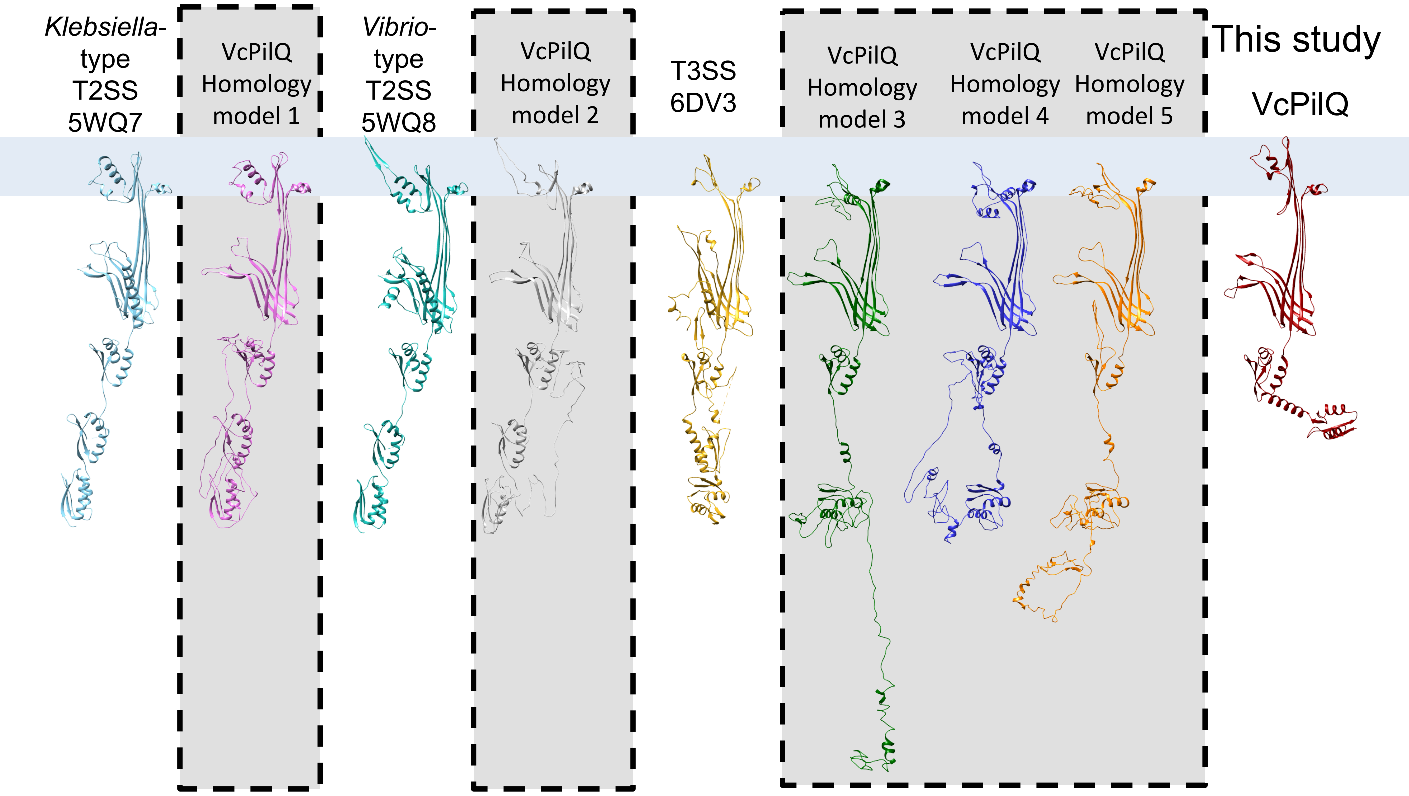

### Supplemental Figure 8

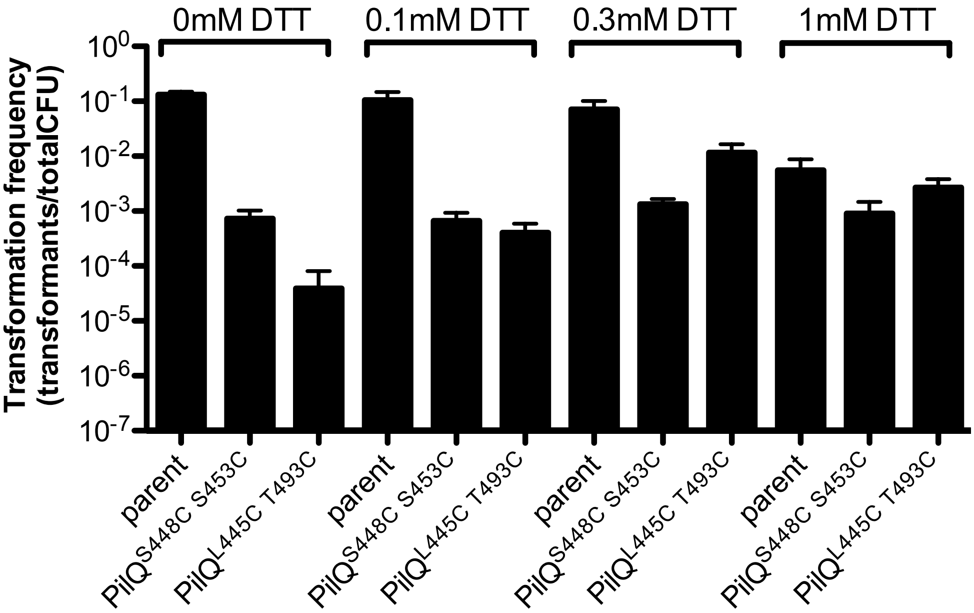

### Supplemental Figure 9

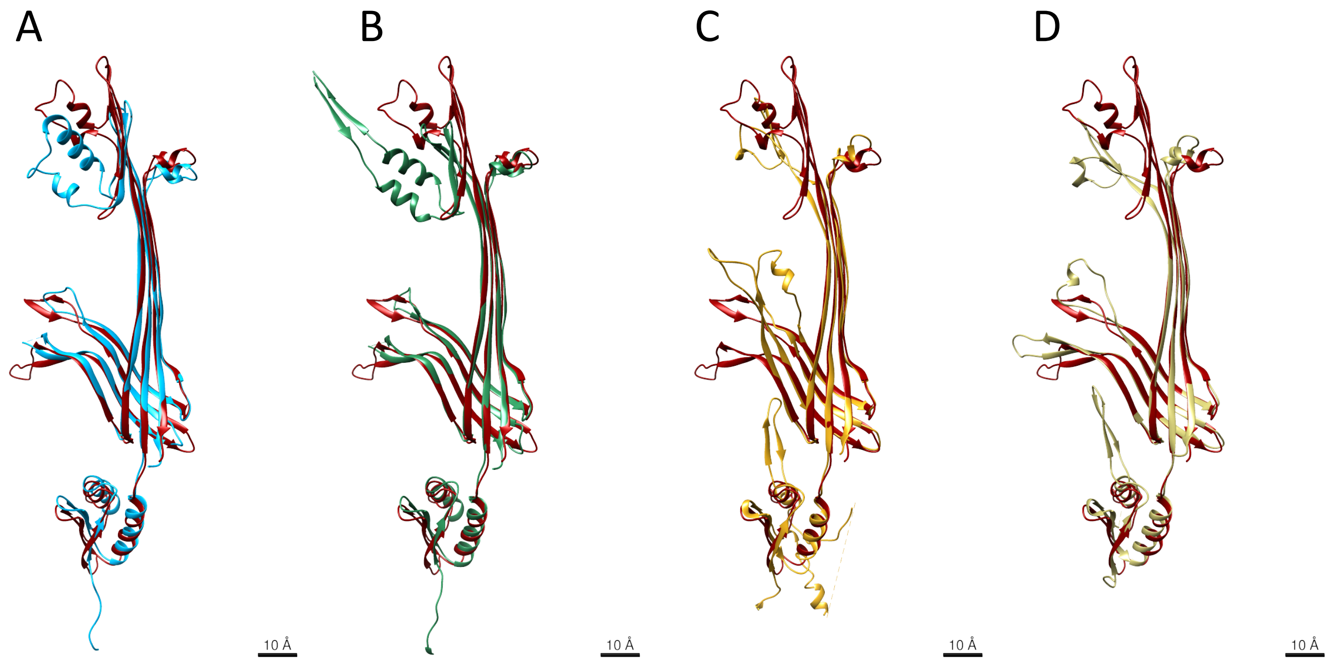


Ec K12 GspD 5WQ7 | Vc GspD 5WQ8 | St InvG 6DV3 | St InvG 5TCQ

E

| VcPilQ Compared to | RMSD (Å) |
|---|---|
| *E. coli* K12 GspD (5WQ7) | 1.2 |
| *V. cholerae* GspD (5WQ8) | 1.2 |
| *S. typhimurium* InvG (6DV3) | 0.97 |
| *S. typhimurium* InvG (5TCQ) | 1.1 |

### Supplemental Figure 10

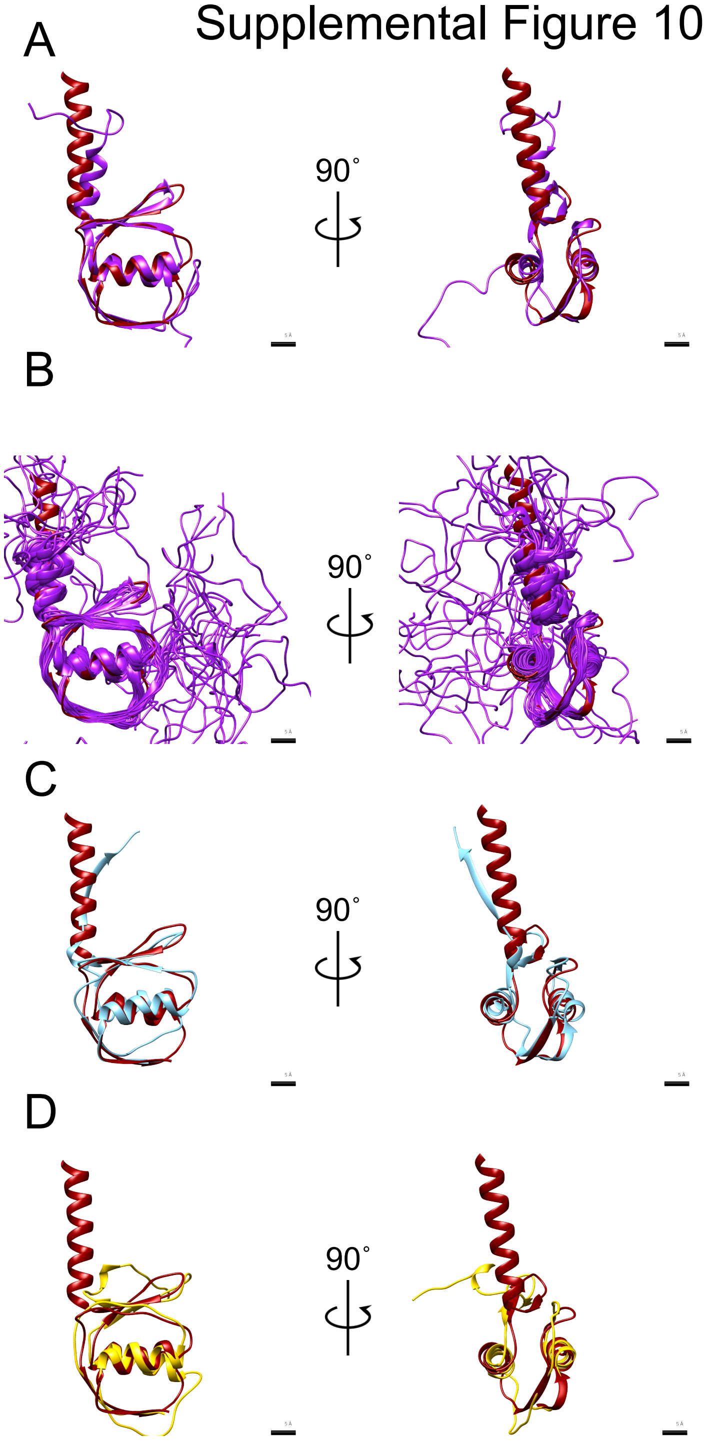


E

| VcPilQ N0 (residues 160-228) compared to | RMSD (Å) |
|---|---|
| *N. meningitidis* PilQ N0 (4AR0, residues 350-417) | 0.9 to 1.2 |
| *S. typhimurium* InvG N0 (6DV3, residues 34-104) | 1.1 |
| *K. oxytoca* PulD N0 (6HCG, residues 27-100) | 1.1 |

#### Supplemental Figure 11

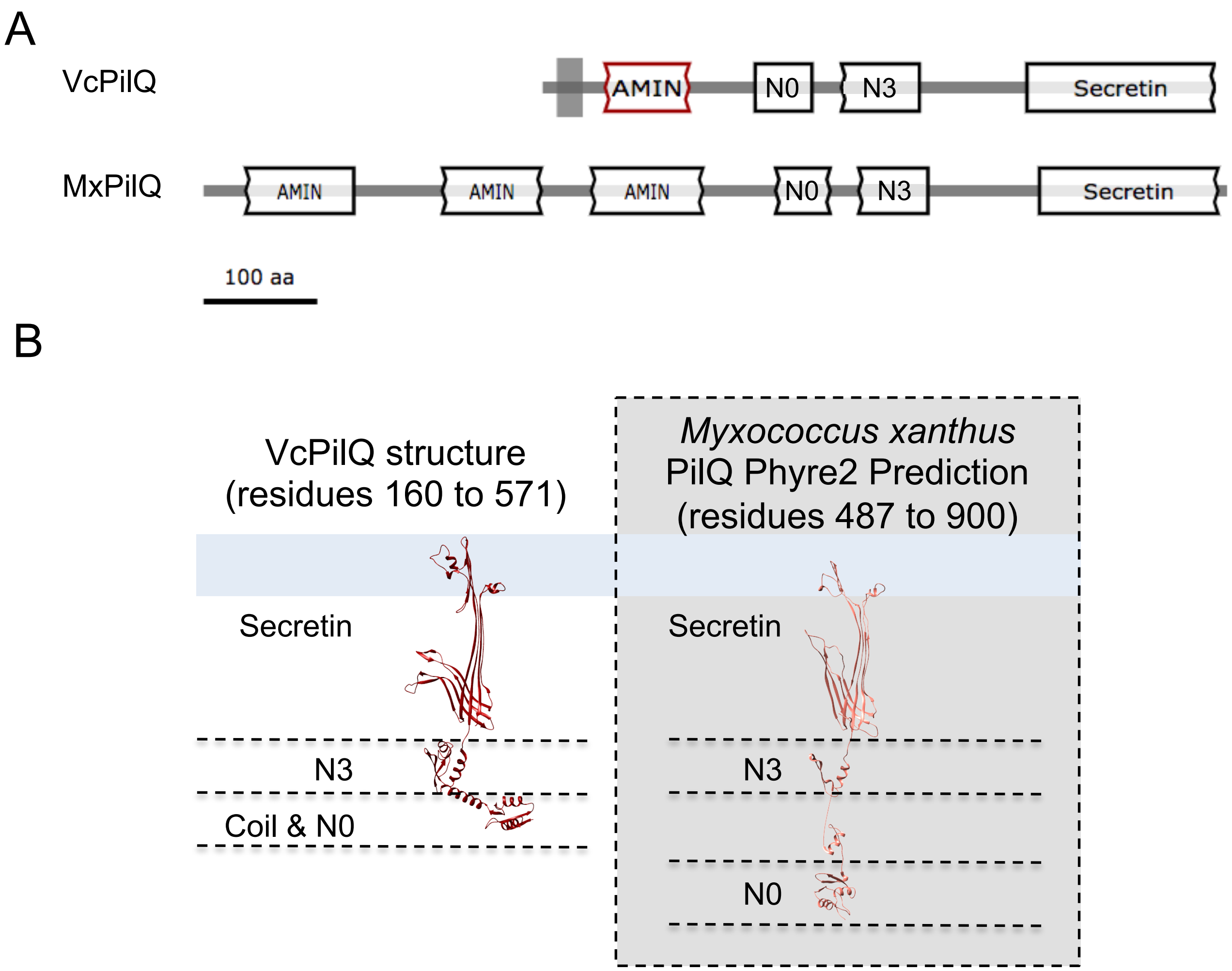
